# Light-induced Trpin/Metout switching during BLUF domain activation in ATP-bound photoactivatable adenylate cyclase OaPAC

**DOI:** 10.1101/2023.09.05.556344

**Authors:** Anaïs Chretien, Marius F. Nagel, Sabine Botha, Raphaël de Wijn, Lea Brings, Katerina Dörner, Huijong Han, Jayanath C. P. Koliyadu, Romain Letrun, Adam Round, Tokushi Sato, Christina Schmidt, Radu-Costin Secareanu, David von Stetten, Mohammad Vakili, Agnieszka Wrona, Richard Bean, Adrian Mancuso, Joachim Schulz, Arwen R. Pearson, Tilman Kottke, Kristina Lorenzen, Robin Schubert

## Abstract

The understanding of signal transduction mechanisms in photoreceptor proteins is essential for elucidating how living organisms respond to light as environmental stimuli. In this study, we investigated the ATP binding, photoactivation and signal transduction process in the photoactivatable adenylate cyclase from *Oscillatoria acuminata* (OaPAC) upon blue light excitation. Structural models with ATP bound in the active site of native OaPAC at cryogenic as well as room temperature are presented. ATP is found in one conformation at cryogenic- and in two conformations at ambient-temperature, and is bound in a non-productive conformation. However, FTIR spectroscopic experiments confirm that the non-productive conformation is the native binding mode in dark state OaPAC and that transition to a productive conformation for ATP turnover only occurs after light activation. A combination of time-resolved crystallography experiments at synchrotron and X-ray Free Electron Lasers sheds light on the initial events around the Flavin Adenine Dinucleotide (FAD) chromophore in the light-sensitive BLUF domain of OaPAC. Initial changes involve the highly conserved amino acids Tyr6, Gln48 and Met92. Crucially, the Gln48 side chain performs a 180° rotation during activation, leading to the stabilization of the FAD chromophore. Cryo-trapping experiments allowed us to investigate a late light-activated state of the reaction and revealed significant conformational changes in the BLUF domain around the FAD chromophore. In particular, a Trp_in_/Met_out_ transition upon illumination is observed for the first time in the BLUF domain and its role in signal transmission via α-helix 3 and 4 in the linker region between sensor and effector domain is discussed.

## Introduction

Light-sensitive photoreceptors evolved in all domains of life, enabling organisms to respond to light variations in their environment. Photoreceptors include a light sensor domain, examples for blue light sensitivity are Light, Oxygen, or Voltage (LOV) domains or Blue Light sensors Using FAD (BLUF) domains^1^. Both offer interesting applications in optogenetics and synthetic biology^2–15^.

BLUF photoreceptors can be found in both bacteria and in eukaryotes^1,16^. They were first discovered in the unicellular flagellate *Euglena gracilis*, in which BLUF regulates the photoactivated adenylate cyclase (PAC) activity^17^. Later, two smaller multi domain BLUF photoreceptors were found in *Beggiatoa sp*. (bPAC)^11,18,19^ and in *Oscillatoria acuminata* (OaPAC)^20–22^, in which they control the enzymatic activity of the linked adenylate cyclase (AC) effector domains. Sequence identity between bPAC and OaPAC is around 58% and the published crystal structure of bPAC (PDB 5M2A) by Linder *et al*.^18^ shares the same overall secondary structure with OaPAC. All BLUF domains exhibit a similar overall structure with an alpha/beta-sandwich ferredoxin-like fold containing a five-stranded β-sheet and two antiparallel α-helices (βαββαββ topology). The flavin-based chromophore binds between the two α-helices on top of the β-sheet^1^.

Due to the small size of 350 or 366 amino acids and their low dark state activity, these two photoreceptors have generated an increased interest in understanding their mechanism in order to use them as optogenetic tools.

PAC from the cyanobacterium *Oscillatoria acuminata* is a 366 amino acid long protein that forms a homodimer containing an N-terminal BLUF sensor domain coupled to an adenylate cyclase (AC) effector domain, the latter being involved in the conversion of ATP into cyclic-AMP (cAMP). cAMP is an important second messenger and plays a crucial role in diverse signaling pathways, such as the mediation of various responses via binding to proteins of the cAMP-receptor protein family in bacteria^7,23,24^.

The previously reported crystal structures of OaPAC (PDB 4YUS, 4YUT) disclosed the conformation of the flavin binding pocket of the BLUF domain involving a conserved tyrosine (Tyr6), glutamine (Gln48) and methionine (Met92) in its Met_in_ conformation^20^. Previous studies have shown the important role of the tyrosine and the glutamine to initiate the photocycle after excitation of oxidized FAD and the signal transmission in the BLUF domain^1,6,21,25–37^. Accordingly, flavin and tyrosine radicals have been identified as intermediates, which then yield a glutamine tautomer and oxidized FAD in the light state. The semi-conserved tryptophan (Trp90) is present for OaPAC in the Trp_out_/Met_in_ conformation in its dark state, which is similar to other BLUF photoreceptors such as bPAC (PDB 5M2A)^18^, TePixD /TII0O7849 (PDB 1X0P)^38^, BlrP1 (PDB 3GFZ)^30^, BlrB (PDB 2BYC)^39^ and BlsA (PDB 6W6Z)^40^. In OaPAC, the α-helix 3 on the opposite face of the FAD binding pocket of the BLUF domain interacts with its symmetry mate in the dimer interface to form a coiled coil.

After comparing the structure of OaPAC in its dark and light state, Okhi *et al*. suggested that the C*γ*− C*δ* bond of glutamine (Gln48) rotates approximately 40°, changing the hydrogen bond network around the isoalloxazine ring and especially between the FAD O4 and the Nε2 atom of Gln48, which could involve a glutamine tautomerization^21^. Mutational studies also suggested that the α-helix 3 bundle is related to the signal transmission from the BLUF domain to the AC domain in order to control the active site^20^.

In the proposed mechanism of bPAC from Linder *et al*., the conformation of the tyrosine and glutamine residues are linked to a kink in β-sheet 4^18^. They also proposed that the loop between β-sheet 9 and β-sheet 10, called the tongue region, is at a central position to act as a lever that facilitates the opening of the AC domains for both subunits. With these, the active site, located at the interface between both AC domains, opens up to enable ATP turnover.

There is an ongoing debate about the presence of a Trp_in_ conformation in some of the BLUF photoreceptor structures^32,33,41^ and the importance of Met_in/out_ and Trp_in/out_ conformations in the signal transmission^30,42–44^, but to our knowledge none of the crystallography studies have yet shown a Trp/Met switch after photoactivation in PAC photoreceptors^18,21^.

A study on the C-terminal region of OaPAC suggested that the C-terminal region regulates the adenylate cyclase activity, hence inhibiting cAMP production^22^. Therefore, the deletion of the C-terminal region of OaPAC increases enzymatic activity and enables higher sensitivity to light.

Despite the large number of studies on the light excitation in OaPAC discussed above, the signal transmission to the AC domain and the overall regulatory mechanism remain elusive. Our study, combining time-resolved crystallography with FTIR spectroscopic analysis, provides information on the mechanism after excitation of OaPAC including the first steps of the signal transmission and allosteric communication of BLUF and AC domain. We demonstrate that photon absorption by the FAD upon photoexcitation leads to a change in the hydrogen bonding network between the conserved tyrosine, the strictly conserved glutamine and the FAD chromophore. This causes the glutamine to rotate 180° and the methionine to move into a Met_out_ conformation. The semi-conserved tryptophan is therefore able to adopt a Trp_in_ conformation, enabling a straightening of the β-strand 5 and further transmitting the signal to the central coiled coil α-helix 3 and the α-helix 4 handle. Our crystallographic data also allow us to observe the presence of native ATP in the OaPAC active site, bound initially in a native but non-productive conformation.

## Results and discussion

### Binding coordination of ATP in the OaPAC active site

In order to unveil the details of native ATP-binding in OaPAC, crystals of OaPAC in its holo-form and without ATP were analyzed with X-ray crystallography.

OaPAC was grown in an orthorhombic crystal form in the dark. A structural model of OaPAC without ATP, called ATP-free OaPAC from here on, could be solved at 1.5 Å resolution at cryogenic temperature and a visual representation of the model is shown in Supplementary Figure S1. The ATP-free OaPAC structure is significantly different to the structures solved by Okhi *et al*., where the protein crystallized either in a hexagonal space group (PDB 4YUS) with 2.2 Å Cα RMSD, or in an orthorhombic space group (PDB 4YUT) with 2.4 Å Cα RMSD, although the overall secondary structure is comparable (Supplementary Figure S1)^20^. OaPAC co-crystallized with ATP in the dark led to a structural model with ATP bound in the active site solved at 2.1 Å (Figure 1A). As the OaPAC dimer interface forms two active site pockets, two ATP molecules are bound to the holoenzyme. An F_o_-F_c_ Polder map contoured at 3 σ reveals binding of ATP in the active site (Figure 1B). This finding represents the first structure of native ATP bound to a wild type photoactivatable adenylate cyclase, as so far ATP has only been observed in a mutated, constitutively active version of bPAC^18^. A comparison between both ATP-free and holo OaPAC reveals which side chains around the active site move upon ATP binding and are involved in ATP coordination (Figure 1C). Notably, only one magnesium ion Mg_A_ is bound in the ATP-free OaPAC structure, while two magnesium ions are involved in ATP coordination in the holo structure (Mg_A_ and Mg_B_).

**Figure 1:**
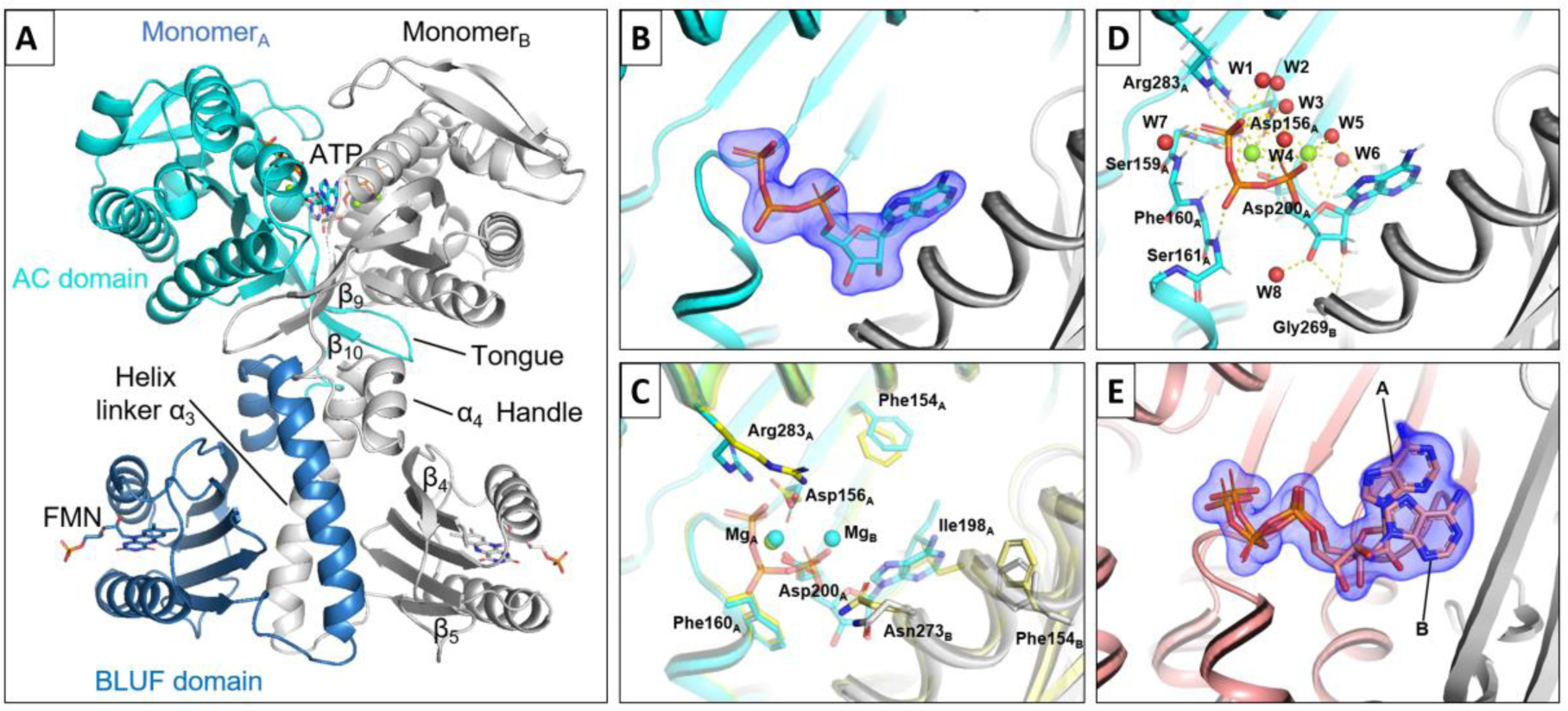
Dark-state structural model of holo OaPAC and details of ATP coordination in the active site. A) Visual representation of holo OaPAC with BLUF domain (blue), AC domain (cyan) of one OaPAC monomer and the second monomer in grey. To each monomer, one FMN is bound in the BLUF domain and one ATP and two magnesium ions are bound to the AC domain. B) ATP binding in a single conformation at cryogenic temperature is visible without bias, as shown by the F_o_-F_c_ Polder map contoured at 3σ. C) Side chain movements upon ATP binding at cryogenic temperature (green=ATP-free, cyan/grey=holo). The movements mainly involve Asp156_A_, Ile198_A_, Aps200_A_, Arg283_A_, Phe154_B_, Asn273_B_. Additionally, two magnesium ions are involved in the coordination of ATP, while only one magnesium ion (Mg_A_) is present in the ATP-free state. D) Detailed view on the ATP coordination at cryogenic temperature via hydrogen bonds by 2 magnesium ions (Mg_A_ and Mg_B_), water molecules, Asp156_A_, Ile157_A_, Ser159_A_, Phe160_A_, Ser161_A_, Asp200_A_, Arg283_A_ and Gly269_B_. The ribose and adenine are flipped by 180° around the adenine N1-N9 axis, revealing that ATP binds in non-productive conformation in the dark. E) ATP binding in two different conformations at room temperature is visible without bias, as shown by the F_o_-F_c_ Polder map contoured at 3σ (red=monomer_A_, grey=monomer_B_). Both conformations have approximately 50% occupancy and they differ by the orientation of their ribose and the position of the adenine.

Asp156_A_ rotates around 80° and interacts with Mg_A_ and W3 in order to enable ATP binding coordination in cooperation with Asp200_A_. The latter, which is involved in the coordination of Mg_A_ in both structures, also rotates 30° to coordinate with the second Mg_B_ ion, not present in the ATP-free OaPAC structure. Ile198_A_ is strongly displaced after 180° rotation of its side chain to enlarge the hydrophobic pocket for the adenosine in the activate site. The Phe154_B_ side chain moves closer to the adenine moiety (3.6 Å in holo vs 3.9 Å in ATP-free OaPAC). Asn273_B_ side chain rotates 180° around Cβ and by this Cγ moves 2.1 Å away to give space to the adenosine. Arg283_A_ turns 4.2 Å away from the surface and Arg283_A_ NH2 forms a hydrogen bond to ATP O1γ (2.9 Å) to coordinate the γ-phosphate of ATP (Figure 1D and Supplementary Figure S2A). Between β-sheet 6 and α-helix 5, several amino acids are involved in binding the phosphates directly, in addition to the coordination via Asp156_A_, Asp200_A_, Ile157_A_ and the two magnesium ions. Ser159_A_ N, Phe160_A_ N1 and Ser161_A_ N1 form hydrogen bonds with ATP O3γ, O2β and O1β, respectively (2.9 Å, 2.7 Å and 2.7 Å). The water molecules W1,3,4,5 and 7 also interact with the phosphates, forming hydrogen bonds to stabilize them. Gly269_B_ N forms a hydrogen bond with O2’ from the ribose (3.1 Å). Notably, the adenine is not coordinated by hydrogen bond with any other residues, but has mainly hydrophobic interactions with surrounding amino acids (Supplementary Figure S2A). W6 and W8 help with stabilization of the ribose. A water molecule near the Mg_B_, which could correspond to W6 in our model, was already described as essential for the proton transfer route during ATP turnover^45,46^. In previous studies from Lindner *et al*., it was shown that the side chain of Phe155 in bPAC mutant (bPAC_Y7F_) moves away to give space for ATP binding, while Glu157 (corresponding to Aps156 in OaPAC) rotates upon ATP binding to facilitate ATP coordination via Mg_B47_.

Grigorenko *et al*. described the most favorable ATP configuration in the AC active site based on the lowest energy barrier to initiate ATP turnover^45^. In their structure, the O3’ from the ribose is oriented towards one of the magnesium ions and is located around 3 Å away from the phosphate Pα, which enables an in-line nucleophilic attack. This conformation facilitates the proton translocation to the neighboring aspartate residue (Asp200), which corresponds to the first step of the enzymatic reaction. However, the binding of ATP substrate in our cryogenic structure differs from the one proposed by Grigorenko *et al*., as the distance between O3’ from the ribose ring and ATP α-phosphate is too large in the presented structure (more than 5 Å away) to permit a nucleophilic O3′ attack on Pα. In our structure, the α-phosphate is rotated away from the ribose ring and the ribosyl O3’ is not coordinated by Mg_B_ but is facing the opposite direction. W6, coordinated by Mg_B_, is therefore not able to interact with the ATP O3’ to facilitate the proton transfer route^45^. Our ATP conformation corresponds to a higher energy state and is not suitable for a low energy proton translocation. Therefore, ATP is bound in our structure in a non-productive conformation and requires additional structural conformational changes before the conversion to cAMP can happen.

After revealing the conformation of ATP in the active site at cryogenic temperature, analysis of the binding at room-temperature was performed to discover potential differences. A near radiation damage-free structure of OaPAC at room temperature in its dark state was determined using serial femtosecond crystallography (SFX) at the European X-ray Free Electron Laser (EuXFEL). The obtained structure is similar to the structure at cryogenic temperature with 0.40 Å Cα RMSD over 350 residues. The resolution is slightly better at room-temperature with 1.8 Å. The unit cell parameters also change from a = 54.5 Å, b = 146.4 Å, c = 104.9 Å at cryogenic temperature to a = 54.3 Å, b = 145.8 Å, c = 105.3 Å for the SFX data. The SFX structure revealed the presence of a second ATP conformation at room temperature with 50% occupancy (annotated A and B) (Figure 1E). In the second conformation B, the phosphates are only displaced 0.3 Å in comparison to the conformation A, but the ribose adopts a different conformation with a torsion angle Pα-O5’-C5’-C4’ of -164° vs. -144°. C5’ C4’ and C3’ are moved 0.5 Å, 1.2 Å and 1.4 Å, respectively, hence coming closer to Gln269_B_ and Asn273_B_. The O3’-Pα distance slightly increases from 5.4 Å in conformation A to 5.5 Å in conformation B. The adenine is dragged away by approximately 5 Å, changing the torsion angle C8-N9-C1’-O4’ from 45° to 77°.

Notably, the second ATP conformation does not seem to affect the structural conformation of the surrounding amino acids in the active site significantly. When comparing the single ATP conformation of our cryogenic structure to the two ATP conformations determined at room temperature, only few noticeable changes are apparent (Supplementary Figure S2A-C). We can observe the loss of a hydrogen bond between Cys201_A_ and Mg_B_, which interacts in the room-temperature structure with a water Asp156_A_ instead, and the formation of hydrogen bonds between Asn273_B_ and ATP. Asn273_B_ is modelled with two conformations in our room temperature-structure and its Cγ atom is dislocated 0.6 Å or 2.1 Å away from the position at cryogenic temperature, respectively. This enables Asn273_B_ to get closer to the ATP and to form hydrogen bonds between Asn273_B_ Nδ2 and ATP O1α for both conformations (3.0 Å for ATP conformation A and 2.7 Å for ATP conformation B). Notably, Asp156_A_ is rotated 70° at room temperature compared to OaPAC with ATP at cryogenic temperature. This enables Asp156_A_ to interact with both magnesium ions at room-temperature, while Asp156_A_ was interacting with MgA and W3 at cryogenic temperature.

Comparing both ATP conformations at room-temperature, we observe that the hydrogen bond network is similar, with changes mainly in the hydrogen bond lengths (Supplementary figure S2B and C). However, we do notice a few changes in interaction distances in ATP conformation B, due to the movement of the ribose closer to monomer_B_. The distance between ATP O2’ and Gly269_B_ is shortened from 3.8 Å to 2.4 Å, which results in formation of a hydrogen bond between them. As the adenine is mainly coordinated by hydrophobic interactions, the 5 Å displacement does not affect the surrounding residues. We do however notice additional hydrophobic interactions between the adenosine moiety of ATP conformation B and monomer_B_ (Ile198_B_, Met203_B_, Ile267_B_ and Asp270_B_), while other hydrophobic interactions (Phe154_B_ and Ala278_B_), present in conformation A and in the structure at cryogenic temperature, have disappeared.

The ATP has a certain amount of room for movement in the active site, as the adenine can move freely within a 5 Å range of motion. However, both conformations found in the room-temperature structure are non-favorable conformations for the production of cAMP, as described by Grigorenko *et al*., and the binding of the most favorable ATP-conformation is not possible in the active site configuration observed in our structures^45^. As our room-temperature data also shows that the adenosine moiety is not strongly coordinated in the active site (Supplementary Figure S2B and C), it is plausible that rearrangements can occur after the AC domain opens up upon light activation.

In order to investigate if ATP can be catalytically turned over to cAMP by the studied OaPAC enzyme upon blue light excitation in solution, time-resolved light-minus-dark FTIR difference spectra were recorded for 8 minutes after excitation by an LED (Figure 2A). Besides contributions from FAD and the enzyme, characteristic negative difference signals from ATP are visible in the spectral range of 900 – 1300 cm^-1^ ^48^. Positive difference signals after blue light excitation match well with the additive signature of cAMP, pyrophosphate and phosphate in solution (Supplementary Figure S3), demonstrating that ATP is indeed turned over to cAMP by the enzyme after exposure to blue light (Figure 2B). As ATP is bound in a non-productive conformation in our crystals, it could be argued that this conformation is an artefact of the crystal packing. To test this hypothesis, FTIR difference spectra were recorded in the absence of ATP and showed that all main difference peaks attributed to the cyclase domain are lost when ATP is not bound, but are mostly restored in the presence of non-hydrolysable ApCpp (Supplementary Figure S4). Accordingly, it appears that the AC domain does not undergo large conformational rearrangements upon blue light excitation when ATP is not present. This leads us to the conclusion that ATP is binding to OaPAC in a non-productive conformation in the dark, while the cyclase domains of both monomers are closed. Our data indicate that blue light induces a conformational change along with the opening up of the AC domains only when ATP is bound, allowing the ribose and adenine moiety of ATP to rotate into a productive conformation prior to pyrophosphate cleavage and cAMP formation. Similarly, no light-dependent increase in ATP affinity was observed in bPAC_Y7F_, where ATP is also bound in a non-productive conformation^18^.

**Figure 2:**
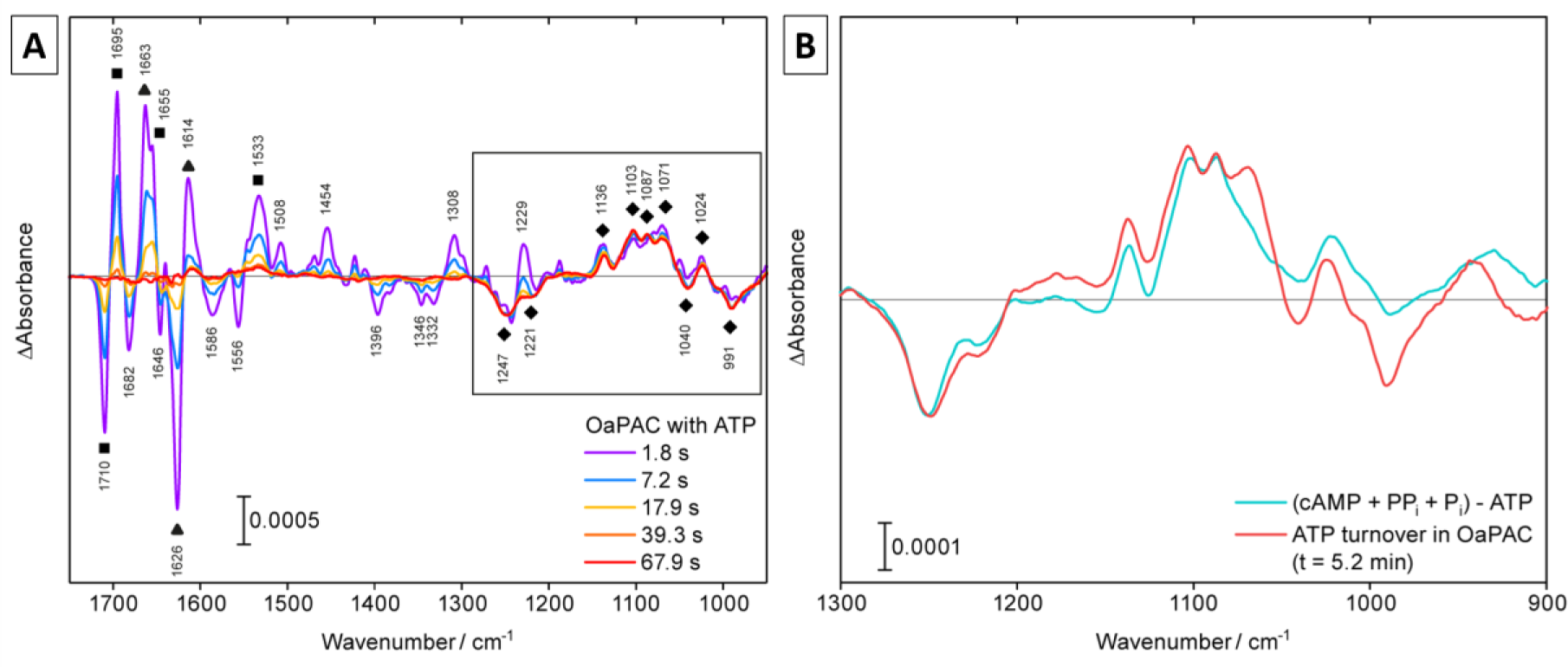
Light-minus-dark FTIR difference spectra of OaPAC in solution in the presence of ATP. A) Signal evolution is observed by time-resolved FTIR spectroscopy. Assignment of contributions by the BLUF and AC domain of OaPAC to the FTIR difference signal are shown, as well as the signals attributed to ATP turnover. Specific difference bands are labelled with squares (BLUF), triangles (AC) and diamonds (ATP), other bands represent mixed contributions. B) Recorded difference signal from ATP matches the expected signal of cAMP, pyrophosphate (PP_i_) and phosphate (P_i_) minus ATP, showing the turnover of ATP to cAMP upon blue light excitation in the FTIR difference signal.

To dissect the contributions to the time-resolved FTIR difference signals of OaPAC by the BLUF domain and the AC domain, a truncated mutant was analyzed that contains the BLUF domain including the long α-helix 3 linker and the α-helix 4 (OaPAC_1-137_) (Supplementary Figure S5). Difference bands were accordingly assigned and labeled with squares for BLUF, triangles for AC and diamonds for ATP turnover (Figure 2A). Other signals contained mixed contributions. The difference spectra show the characteristic 1710(-) and 1695(+) cm^-1^ carbonyl signature of BLUF, which was reported for the change in hydrogen bonds to C4=O of the flavin^49–51^.

Some prominent bands originate from the AC domain as they are still present in the double difference spectrum after subtracting the OaPAC_1-137_ contribution from the full-length protein (Supplementary Figure S5). Accordingly, the 1663(+) cm^-1^ band was determined to originate from the OaPAC cyclase and might correspond to the α-helical signal at around 1650(+) cm^-1^ in bPAC^49^. It shows the same characteristic peak shape with a low frequency shoulder. Interestingly, this shoulder was attributed to the cyclase in bPAC and was lacking in the bPAC-BLUF_1-124_ domain spectrum without α-helix 4 linker, whereas here a positive signal at 1655(+) cm^-1^ is observed for OaPAC_1-137_ containing the α-helix 4 linker. Accordingly, we assign this signal at 1655 cm^-1^ to a response in helical structure caused by the presence of the α-helix linker 4 linking BLUF and cyclase, which has been suggested to be important for signal transfer in BLUF photoreceptors^30,52–55^.

Large difference bands from the AC domain are also present at 1626(-) and 1614(+) cm^-1^ (Supplementary Figure S5). In bPAC, the corresponding negative signal was found at 1631(-) cm^-1^ and could be attributed by its characteristic frequency to a β-sheet response in the cyclase domain ^49^. This signal is unlikely to originate from a suppressed β-sheet 5 response in the BLUF domain because signaling from FAD to α-helix 3 and α-helix-4 via β-sheet 5 is intact in OaPAC_1-137_. The positive band at 1614 cm^-1^ has not been observed in bPAC^49^ and YcgF^56^ and is very low in frequency for an amide I signal of secondary structure elements^57^. Steady-state measurements on OaPAC and OaPAC_1-141_ under deuterated conditions^58^ did not show any prominent positive bands in this region either, which might be explained by a downshift to 1560 cm^-1^ or by band compensation effects (Supplementary Figure S5).

After assigning the contributions of both domains to the recorded difference bands, time constants for the relaxation of structural changes in the BLUF as well as the AC domain could be derived from the evolution of the FTIR difference spectra over time. For the BLUF domain in OaPAC, the signals at 1710 (-)/ 1696 (+) cm^-1^ decay with τ = 8.9 ± 0.5 s. Of note, the relaxation was significantly faster in the OaPAC_1-137_ mutant (τ = 6.3 ± 0.2 s) and gave similar results as full-length OaPAC in the absence of ATP (τ = 5.8 ± 0.7 s) (Figure 3). This comparison indicates that the binding of ATP and turnover to cAMP has an allosteric feedback effect on the relaxation of BLUF and its lifetime. The relaxation of the AC domain was investigated using the 1615 cm^-1^ band and yielded τ = 2.9 ± 0.6 s in the presence of ATP (Figure 3), much faster than the signals of the BLUF domain. As all signals from the AC domain are lost in the FTIR spectra in the absence of ATP (Supplementary Figure 4), meaning that the AC domain is not activated without ATP bound, a relaxation without ATP could not be determined.

**Figure 3:**
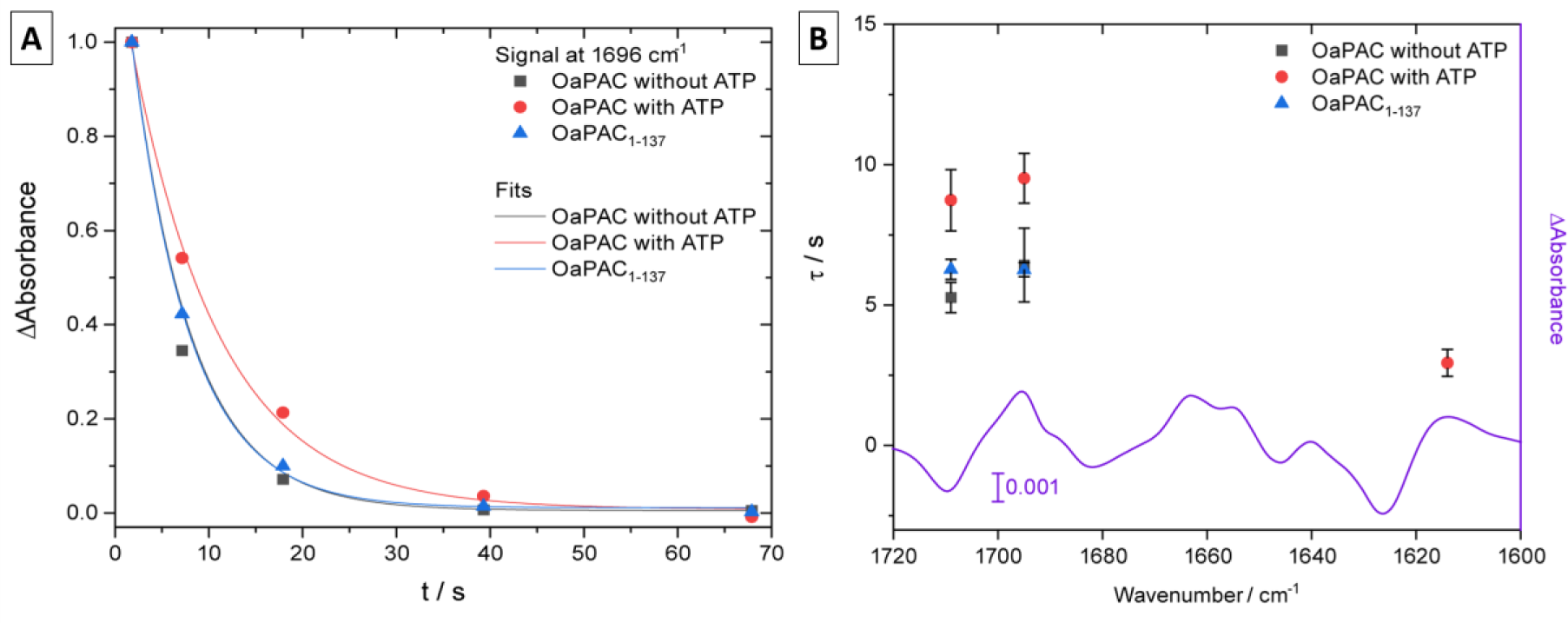
Kinetics of relaxation of structural changes after excitation of OaPAC derived from FTIR difference signals. A) The evolution of the time-resolved FTIR difference absorbance was plotted at 1696 cm_-1_ for OaPAC in the presence and absence of ATP and for OaPAC_1-137_ and analyzed by monoexponential functions. B) The resulting time constants for relaxation of the BLUF domain at 1710 and 1696 cm_-1_ and for the AC domain at 1615 cm_-1_ are summarized. The AC domain shows a significantly faster relaxation (τ = 2.9 ± 0.6 s) than the BLUF domain (τ = 8.9 ± 0.5 s). Relaxation of BLUF becomes faster when ATP is absent or in the truncated OaPAC_1-137_ pointing to a tight allosteric communication.

### Effects of photoactivation in the surrounding of FAD in the BLUF domain in OaPAC

To better understand the reaction of BLUF photoreceptors upon photoactivation and to shed light on the differing opinions in the scientific community about the early steps of this process, a time-resolved SFX (TR-SFX) experiment was conducted at the SPB/SFX instrument of the EuXFEL. Our TR-SFX study describes structural changes happening in the BLUF photoreceptor OaPAC when crystallized with ATP in the orthorhombic crystal form at two pump-probe delays (1.8 µs and 2.3 µs). Shorter pump-probe delays up to 500 ns have been investigated as well, but did not result in interpretable difference signal and are therefore not presented.

Fourier difference density maps for these pump-probe delays calculated against the dark data set using Xtrapol8^59^ show significant differences in the vicinity of the flavin chromophore (Figure 4A, D). The FAD molecule bound in the BLUF domain was modeled as flavin mononucleotide (FMN), as the AMP moiety was not visible in the electron density. Xtrapol8 was used to determine occupancies of the light-excited state and thereby extrapolate structure-factor amplitudes, allowing us to model the structural changes happening at these three time points. The light-excited states after 1.8 and 2.3 µs were estimated at 25% occupancy for both time point. The Fourier difference density maps provide clear evidence of the involvement of the conserved amino acids Tyr6, Gln48 and Met92 in the early steps of the photoactivation (Figure 4A, D).

**Figure 4:**
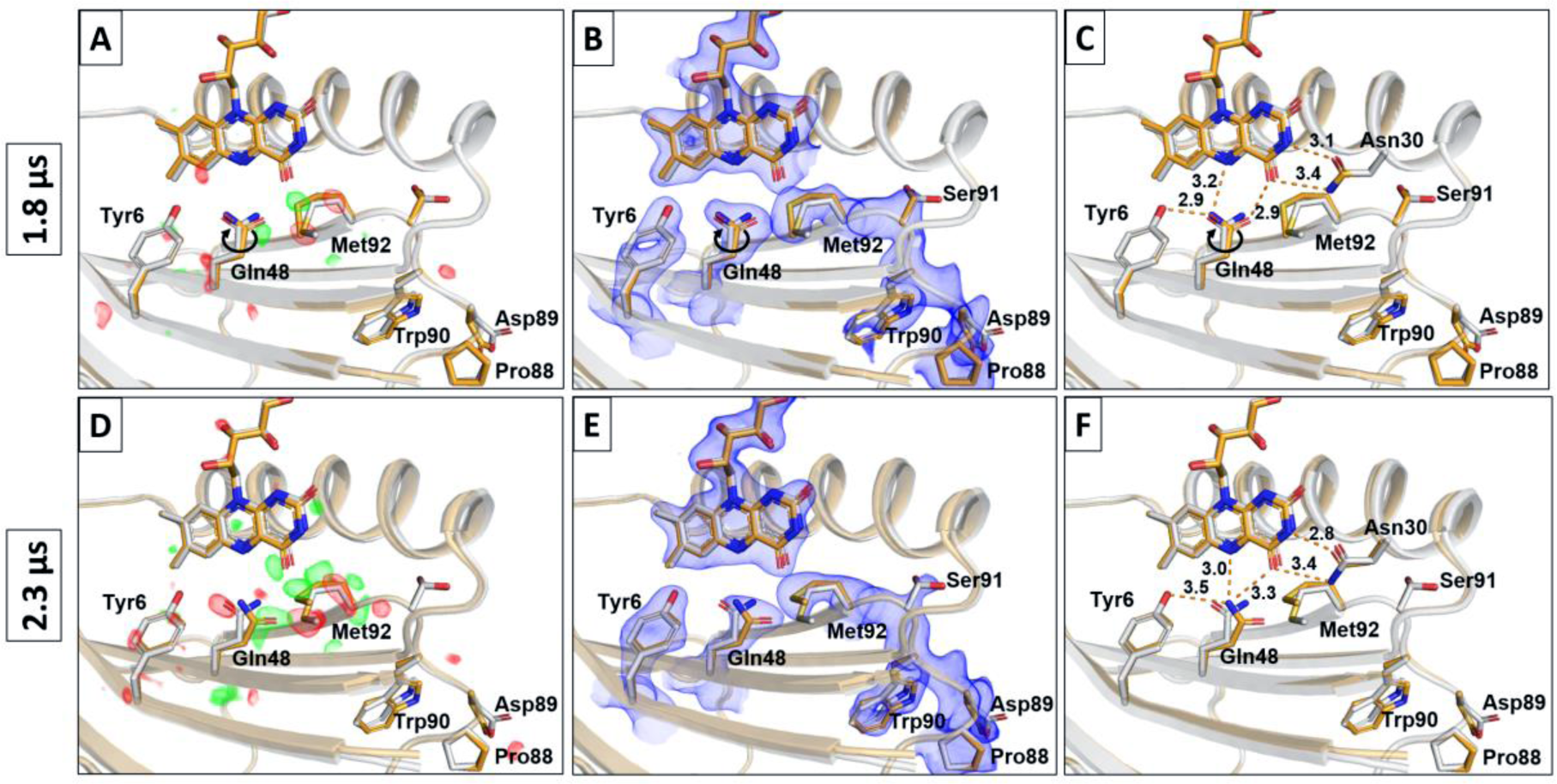
BLUF activation via blue light excitation of flavin chromophore in OaPAC. Structural models of excited OaPAC (orange) with pump-probe delays of 1.8 µs (A-C) and 2.3 µs (D-F) are shown and involved amino acids in stick representation along with the dark model (grey). A and D) F_o-light_-F_o-dark_ difference maps contoured at +3 σ (green) and -3 σ (red) reveal initial changes around the flavin chromophore after excitation. B and E) Extrapolated Fourier difference density maps (2F_o_-F_c_) contoured at 1 σ for the different pump-probe delays, calculated against the dark data set using Xtrapol8 to allow building of the structural model of the excited state. C and) Hydrogen bond distances in Å around the flavin chromophore upon BLUF excitation at the investigated pump-probe delays.

Firstly, after a pump probe delay of 1.8 µs in our experiment, several changes arise, involving the residues mentioned previously (Tyr6, Gln48 and Met92) (Figure 4A-C). It is well known that these H-bonding residues around the flavin are highly conserved among BLUF photoreceptors^21,25–33,35^, making them the likely candidates as functional residues for the early steps of photoactivation. The most significant change concerns Gln48, which has undergone a full 180° rotation, rearranging completely the hydrogen bond network in the chromophore pocket. This rotation breaks the hydrogen bond between FMN O4 and Gln48 Nε2, as the latter is now interacting with Tyr6 OH and FMN N5. The distance between Gln48 Nε2 and Tyr6 OH is increased to 2.9 Å compared to 2.6 Å in the dark state between Gln48 Oε1 and Tyr6 OH (Figure C and Supplementary Figure S7B). Gln48 Oε1 moves closer to FAD O4 (2.9 Å), which creates a new hydrogen bond between them. Gln48 Oε1 also now shows dipole interactions with Met92 Cε being 3.3 Å away and an intramolecular interaction with Met92 Sγ of 3.1 Å, whereas the Gln48 Nε2–Met92 Cε or Sγ distances were 3.6 Å and 3.4 Å in the dark, respectively. In the meantime, Met92 initiates a movement away from the chromophore binding pocket and is displaced more than 1 Å from its original position, as also previously described by Jung *et al*. for AppA^60^. More precisely, the conformation of the Met92 changed with Cε being rotated 90° down and now facing towards β-sheet 2, compared to the dark state. Asn30 Nδ2 and Oγ1 atoms are also displaced 0.3 Å and 0.4 Å further from the flavin compared to the dark state, respectively.

It has been previously discussed that the first steps upon photoexcitation involve a PCET from the conserved tyrosine (Tyr6) to the FAD for AppA and SyPixD/Srl1694^31,34,35,37,60–62^, inducing a change in the hydrogen bond network surrounding the chromophore. Recently, Collado *et al*. suggested a structural mechanism of signaling in OaPAC from time-resolved infrared spectroscopy data, in which a concerted proton electron transfer and the formation of a neutral flavin radical (FADH**·**) as an intermediate species are described^58^. The latter reacts with the tyrosine radical (Tyr6**^•^**) to form the light state of BLUF with oxidized FAD. As the resolution of our crystallographic data is limited to 1.9 Å, it cannot confirm the involvement of proton and electron transfer from Tyr6 to the flavin, but we can confirm the rearrangement of the hydrogen bond network between Tyr6, Gln48, Met92 and flavin in the early steps of photoactivation.

The rotation of Gln48 has also been in constant discussion over the last two decades. Some spectroscopic and theoretical studies show the tautomerization of the glutamine^27,36^, accompanied by its side chain rotation^31,37^, whereas others argue that the tautomerization is not followed by any 180° side chain rotation^21,63^. While it is difficult to assign which tautomer is present with X-ray crystallography data, our TR-SFX study strongly agrees with the presence of a glutamine rotation in the early steps of photoactivation. The rotation of Gln48 was also proposed by Jung *et al*. as the proton-coupled electron transfer (PCET) in AppA creates unfavorable interactions such as the proximity of the protonated FAD N5 to the amide group of Gln48 and of Tyr6 OH to Gln48 Oε1^60^. Therefore, Gln48 rotation restores favorable interactions between each of these groups. Interestingly, this result contradicts the observation made by Okhi *et al*. for OaPAC, where a rotation for Gln48 of only 40° was reported after 20 seconds of illumination (PDB 5X4T)^21^.

2.3 µs after photoexcitation, the changes around the chromophore are more pronounced (Figure 4D) and both Gln48 and Met92 continue their rearrangement (Figure 4D-F). Gln48 undergoes an additional rotation of 40°, which means the rotation is over 220° from its original position in the dark state. It also moved further away from Tyr6, resulting in a distance of 3.5 Å between Gln48 Nε2 and Tyr6 OH. This movement induces Gln48 Oε1 to lose the interaction with FMN O4, and Gln48 Nε2 to interact with both FMN N5 and O4, being 3.0 Å and 3.3 Å away, respectively. Interestingly, a 40° rotation was observed after 20 s illumination in OaPAC by Okhi *et al*., but without the initial 180° rotation of the side chain^21^. This additional 40° rotation is probably necessary to stabilize the hydrogen bonding network around the flavin chromophore and especially with Met92. Indeed, the rotation is coordinated with the displacement of Met92, which flips further away from the chromophore pocket. This movement was clearly visible in the Fourier difference electron density map (Figure 4D) and has been successfully transposed into extrapolated density maps (Figure 4E), allowing us to model Met92 1.5 Å away from its original position. This displacement induces movements in the backbone of adjacent amino acids. All the amino acids in the loop region between β-sheet 4 and 5 (Glu79 to Ser 91) and the following amino acids in β-sheet 5 and the loop between β-sheet 5 and α-helix 3 (Gln93 to Phe103) are destabilized by the movement of Met92. Therefore, the movements of the residues around the chromophore seem to initiate a signal transmission involving more distant amino acids from the chromophore after 2.3 µs in our study.

Notably, the kinetics of these structural changes are significantly slower compared to the protein OaPAC in solution, for which the hydrogen bond rearrangement already happens after a few hundreds of picoseconds^58^. This rearrangement is only visible at 1.8 µs in our TR-SFX study and could not be observed for the faster time points studied up to 500 ns. Slower kinetics have been observed several times in crystallographic studies^64–66^. The difference in dynamics is most likely caused by a different protein environment due to the crystal packing or the crystallization solution^67,68^.

### Signal transduction after photoexcitation in OaPAC

In order to investigate the signal transduction from the BLUF domain to the AC domain after blue light excitation, we performed cryo-trapping experiments to capture a late stage of the reaction. A crystal of OaPAC co-crystallized with ATP was flash cooled immediately after 5 s constant light exposure. By this, full reversion to the dark state could be prevented and X-ray diffraction data of the light-activated sample were collected at 100 K the synchrotron. The structural model of cryo-trapped OaPAC could be solved at 1.7 Å resolution.

Comparison of the structural model between the dark state and the 5 s light activated steady-state, both at cryogenic temperature, showed significant differences in the BLUF domain and especially around the flavin chromophore (Figure 5A-B), the Cα RMSD being 1.36 Å when comparing both dimers and 1.93 Å for an all-atom RMSD. Several main changes around the chromophore are important to notice. The first change involves the hydrogen bond network between the highly conserved amino acids Tyr6 and Gln48 and the flavin (Figure 5B). In dark conditions, Gln48 Nε1 forms a hydrogen bond with FMN O4 (2.9 Å) and FMN N5 (3.3 Å), and Gln48 Oε1 to Tyr6 OH (2.8 Å) (Supplementary Figure S7A). In the cryo-trapped data, the Gln48 side chain undergoes a 205° rotation and the distances of Gln48 Oε1 to FMN O4 is increased to 3.1 Å and from Gln48 Nε1 to FMN N5 to 3.4 Å, and the hydrogen bond between Gln48 Nε1 (replacing Oε1) to Tyr6 OH stretches to 3.1 Å (Figure 5B). The rotation of Gln48 is similar after the continuous 5 s illumination compared to 2.3 µs delay in our TR-SFX. However, the displacement of the Gln48 away from Tyr6 is less pronounced (Gln48 Nε1 being 3.1 Å from Try6 OH after 5 s illumination, vs 3.5 Å after 2.3 µs). This difference is probably due to Gln48 Oε1 forming a new hydrogen bond with Trp90 Nε1 (3.4 Å). This becomes possible as Trp90 moves from the Trp90_out_ conformation to a Trp90_in_ conformation upon light exposure, taking the position of Met92, which is now in a Met92_out_ conformation (Figure 5A and Supplementary Figure S8). The orientation of the glutamine in this model allows a favorable interaction between Gln48 Oε1 and Trp90 Nε1, which would not be the case if Gln48 had not performed a side chain rotation compared to the dark state. The formation and breakage of a hydrogen bond between the carbonyl group of Gln63 and the indole nitrogen of Trp104 in AppA was already proposed by Masuda *et al*. as critical for the light signaling pathway^42,43^. The onset of this Trp_in_ movement can already be seen in our TR-SFX data, where Gln48 has undergone a 180° rotation after 1.8 µs delay and Met92 flips away from the chromophore pocket at the same time, which opens up the space for Trp90 to move in.

**Figure 5:**
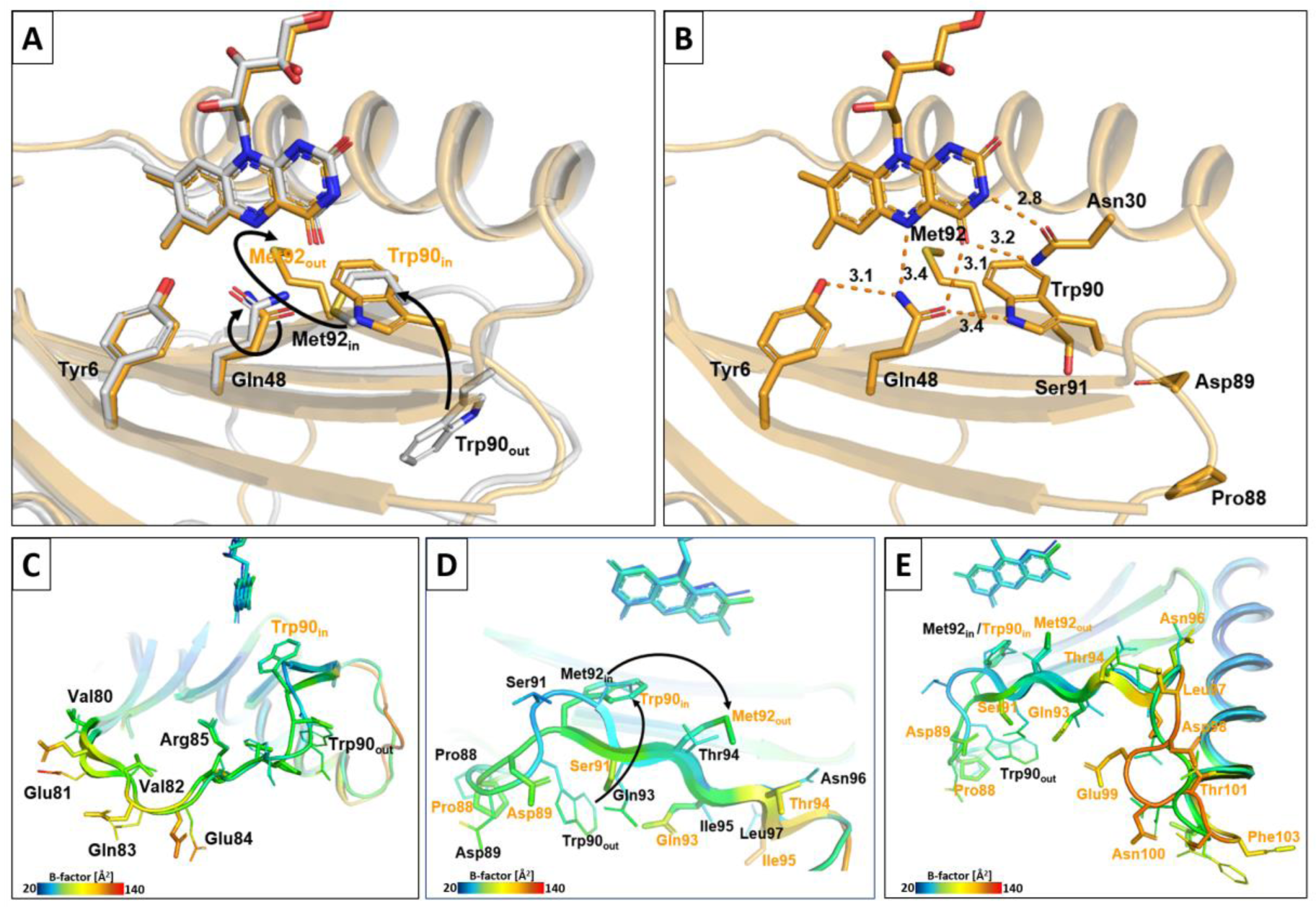
OaPAC shows a transition from Trp90_out_ to Trp90_in_ in cryo-trapped data after 5 seconds of light excitation. A) Structural model of excited OaPAC (orange) is shown and involved amino acids in stick representation along with the dark model (grey). Trp90_out_ to Trp90_in_ transition as well as Met92_in_ to Met92_out_ transition is visualized by arrows. B) Hydrogen bond distances around the flavin chromophore of the Trp90_in_ state. C-E) Trp90_in_ transition is facilitated by mobilization of neighboring amino acids (Val80 to Leu104), as can be seen by the increase in temperature factors. Significantly displaced amino acids in the Trp90_in_ model are labeled in orange. S-pattern shape of β-sheet 5 in the dark structural model straightens to accompany the two-amino acid shift in the Trp90_in_ model (D). This shift is compensated by an increased flexibility in the loop between β-sheet 5 and α-helix 3 (E).

Our cryo-trapped data provide again evidence that the Gln48 rotation is an essential part of the photoactivation process. Further, it sheds light on the Trp_in/out_ and Met_in/out_ conformation, which has been a topic of debate since the first studies on BLUF photoreceptors. So far, it has been reported that some BLUF variants have a Trp90_out_ conformation in the dark-adapted state like OaPAC^20^, bPAC^18^, AppA^60^, SyPixD/Srl1694^33^ in 9 subunits out of 10, TePixD/TII0O78^38^, BlrB^39^, BlrP1^30,69^, BlsA^40^ while others have a Trp90_in_ conformation like SyPixD/Srl1694^33^ in 1 subunit out of 10 and AppA^32,41^. However, it has not been shown that the transition between both conformations occurs upon light exposure as part of the activation mechanism. Consequently, our results contradict the observations made on OaPAC before, where a 20 seconds illuminated structure only showed minor changes upon blue light excitation^21^. More precisely, the rotation of the Gln48 was reported to be only 40°, while we observe a complete rotation of the side chain. Additionally, Met92 only moved away from Gln48 after 20 seconds in the hexagonal crystal form, stretching the distance from 2.6 Å to 3.3 Å, and Trp90 only underwent a 40° rotation. Here, we observed a complete Trp90_in_ conformation in combination with Met92_out_ and a major structural rearrangement of the loop between β-sheet 4 and 5, as well as β-sheet 5 already after 5 s illumination.

It is also interesting to note that a 20 second illumination could not be performed on the orthorhombic crystal form in our study, as the resolution decreased considerably if illuminated for more than 5 seconds, and this even applied when the LED light intensity was significantly reduced. This indicates that a prolonged light exposure disrupts the crystal lattice as OaPAC undergoes its enzymatic activity.

Brust *et al*. suggested the semi-conserved tryptophan to be a key residues in the signal transmission from the FAD chromophore to the protein backbone^29^. Studies on AppA^44^, BlrP1^30^ and BlrB^63^ already proposed the Trp_out_ toward Trp_in_ change in conformation to be one of the first steps in the signal transmission to the effector domain, but it has never previously been described for PAC. So far, structures of OaPAC and bPAC only showed the presence of a Trp_out_ conformation in the light-adapted state^18,21^.

The Met/Trp switch here is facilitated by a restructuring of the loop between β-sheets 4 and 5, as well as β-sheet 5 (Pro88 to Phe103). The section of Pro88 to Gln93, which forms an S-pattern in the dark structure, straightens up in the 5 s cryo-trapped data and allows Trp90 to move in, causing the subsequent amino acids to move by two amino acids towards the C-terminus (Figure 5C and D). Consequently, the β-strand 5 is straightened and B-factors from Gln93 to Leu104 increase to over 100 Å^2^. In particular, the loop between β-sheet 5 and α-helix 3 (Asn96 to Leu104) becomes disordered (Figure 5E). Most likely, Aps98 to Thr101 bulges out and reduces the C-terminal shift to only one amino acid. By position Leu104 at the beginning of α-helix 3, the shift is fully compensated for and all subsequent amino acids align with the positions from the dark structure. FTIR studies by Masuda *et al*. on the AppA Trp104Ala mutant (corresponding to Trp90Ala in OaPAC) revealed that a specific signal, attributed to changes in the β-sheet structure after light activation, was eliminated by the mutation. This confirmed that the tryptophan residue is involved in the transmission of the signal through a specific β-sheet structure modification^42^. Jung *et al*. also speculated that the kink in the β-sheet 5 represents the dark-adapted state of AppA, whereas the regular β-strand corresponds to the light-adapted state^60^. Our observations agree with both proposals as they confirm the implication of Trp90 specifically in the straightening of the β-sheet 5 during the signal transmission in the light-adapted state of OaPAC.

In this study, we present strong evidence that a Met/Trp switch is present in OaPAC upon illumination that is of significant importance in the signaling mechanism. This novel observation enables us to propose Trp90 to be a main actor in the signal transduction after blue light excitation of the chromophore, dragging the entire β-strand 5 into a new conformation and inducing subsequent structural changes for signal transmission. A visual presentation of the changes around the flavin chromophore after light excitation based on the TR-SFX data and the restructuring accompanying the Met/Trp switch based on the 5 s cryo-trapped data are shown in Supplementary Video 1.

However, some questions remain, such as the specific role of the Met/Trp switch in the signal transduction. It could either be functionally relevant as an essential part of the signal transmission to the AC domain, which would take place via the α-helix 3 linking the BLUF domain to the AC effector domain; or it could be only a mechanism contributing to the stabilization of the light activated state and therefore the reason for the long-lived light-activated state of OaPAC.

The semi-conserved tryptophan residue Trp90 has been proposed to be related to the slower dynamics discovered in the signal transmission of AppA and OaPAC^29,70^. In particular, Brust *et al*. studied the relaxation pathway of AppA with vibrational spectroscopy^29^. They discovered a hierarchical pathway, where structural changes associated to H-bonded residues in the flavin binding pocket relax more slowly than changes in more distant residues. They also studied the influence of a Trp104Ala mutant in AppA (Trp90Ala for OaPAC) and revealed that the relaxation to the dark state was considerably accelerated by a factor of 80. The structural changes observed in more remote residues in AppA wild type were suppressed in the mutant, indicating that long ranged structural changes critical for the protein function did not develop without the presence of the tryptophan. These data confirm that Trp90 is a key residue in communicating the signaling state to the protein backbone in AppA and OaPAC. In addition, the importance of the α-helix 3 in signal transmission in OaPAC was proposed by Okhi *et al*., after showing the suppression of the AC activity upon mutation of both Leu111 and Leu115, which form hydrophobic interactions between the α-helices 3 across the dimer interface^20^. They also described the loss of enzymatic activity when mutating Tyr125 (α-helix 3) or Asn256 (β-sheet 5 in the AC domain), as these two residues share an inter-subunit hydrogen bond. These two observations confirm the implication of α-helix 3 in signal transmission, with the necessity of critical non-polar and polar interactions across the dimer interface. These previous studies would support the proposal that Trp90’s function is not only to stabilize OaPAC and enable its long-lived state, but also its involvement in the mechanism contributing to the signal transmission.

Interestingly, for bPAC Linder *et al*. proposed the kink of β-sheet 4 as a pathway for signal transmission from the FAD chromophore to the α-helix 3 to 4 linker region before going through the tongue region of the AC domain^18^. Our data, however, do not confirm this possibility for OaPAC and tend rather toward a signal transmission via the β-sheet 5 to the α-helix 3 linker. Residues in the β-sheet 4 close to the α-helix 3 or α-helix 4 do not show significant change upon light activation, whereas residues in the loop between β-sheet 5 and α-helix 3 exhibit strong movements with a significant increase in B-factors in that region, confirming the involvement of the loop in the signal transmission.

Notably, the observed changes in the cryo-trapped data versus the dark models are limited to conformational changes in the BLUF domain, while only minor changes are observed in the AC domain. Evaluation of the temperature factors on the entire structural model of OaPAC showed that they are consistently higher in the BLUF domain (Supplementary Figure S8). They confirm distinct conformational changes in the entire backbone of β-sheet 4, β-sheet 5 and α-helix 3, including the loops between them. Interestingly, the temperature factors also indicate that the α-helix 4 region is affected during signal transmission. However, the AC domain seems to be particularly stable after 5 s constant illumination and ATP is still clearly visible in the electron density. Since after a long exposure of 5 seconds before cryo-trapping the reaction is expected to progress further towards ATP to cAMP turnover, FTIR light-minus-dark difference spectra were recorded from a crystal slurry in orthorhombic space group. When comparing the spectra with those of full length OaPAC and OaPAC_1-137_ in solution, it can be noted that the difference signal in the crystal packing resembles the OaPAC_1-137_ containing only the BLUF domain and the α-helix 3 and 4 linker region, showing that the AC domain does not undergo large conformational changes in the crystal form (Figure 6A).

**Figure 6:**
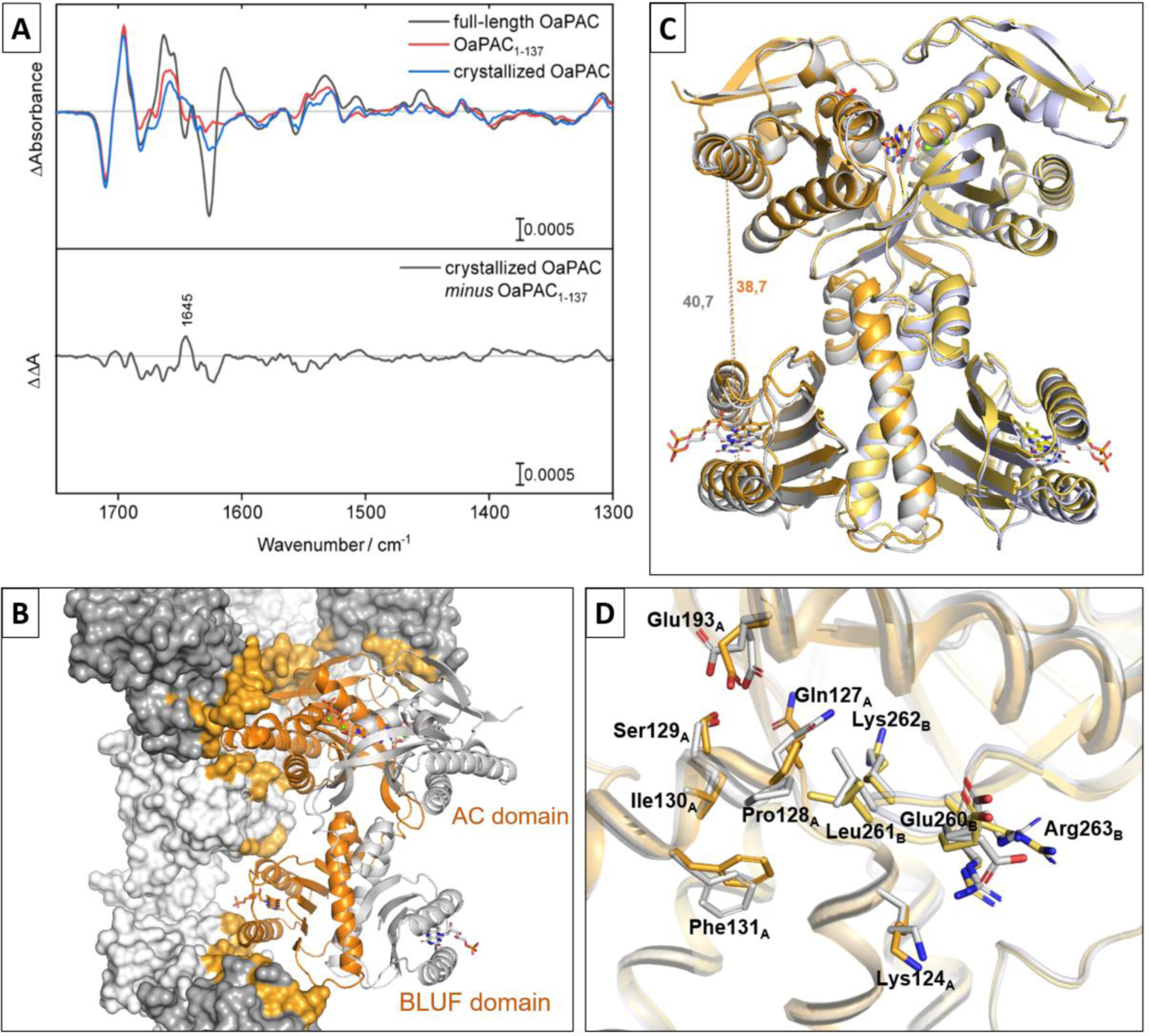
Analysis of adenylate cyclase (AC) domain reaction upon light exposure in orthorhombic OaPAC crystals. A) Comparison of FTIR difference signal after blue-light exposure of OaPAC and OaPAC_1-137_ in solution with OaPAC in orthorhombic crystal packing. Recorded difference signal of crystallized OaPAC matches the OaPAC_1-137_ containing the BLUF domain and α-helix 3 and 4 only, showing that the AC domain is not undergoing large conformational changes in the crystal form. B) Structural model of 5 s cryo-trapped OaPAC (orange) along with neighboring molecules in orthorhombic crystal packing as surface presentation (grey). Crystal contacts in a radius of 4 Å around OaPAC are highlighted in orange surface coloring. While the AC domain is sterically locked in place by the crystal contacts, the BLUF domain is less densely surrounded in the orthorhombic crystal packing, giving it a higher mobility. C) BLUF domain moves about 2 Å closer to the AC domain in 5 s cryo-trapped OaPAC (orange) compared to the dark-state model (grey). D) Side chain rearrangements in the α-helical linker region between BLUF and AC domain in 5 s cryo-trapped OaPAC (orange) compared to the dark-state model (grey).

The lack in response from the AC domain upon photoexcitation in both our crystallography and FTIR data can be explained by the high number of crystal contacts present around the AC domain in the orthorhombic crystal form (Figure 6B). The analysis of the crystal packing reveals that movements of the effector domain are sterically hindered by the molecules surrounding the AC domain in the crystal. Indeed, whereas only a few crystal contacts are present around the BLUF domain, four adjacent molecules keep the AC domain locked in place with only a small margin for movement. Consequently, although the BLUF domain of OaPAC in this crystal form can be excited and conformational changes are observed as part of the activation and signal transmission up until the α-helix 4, movements in the AC domain are hindered by the tight crystal packing.

After understanding the sterically hindered reaction of the AC domain in our orthorhombic crystal form, we were able to consolidate the bigger picture of the observed changes in our cryo-trapped structure. Overall, the BLUF domain moves slightly away from the other BLUF protomer and closer to the AC domain. For example, the distance between Ile22 Cα in α-helix 1 (BLUF domain) and Glu224 Cα in α-helix 7 (AC domain) is reduced from 40.7 Å to 38.7 Å in the cryo-trapped data (Figure 6C). This movement could be with a prelude to the hinge motion of the tongue section between BLUF and AC (loop between β-sheet 9 and 10) proposed by Lindner *et al*. As the opening up of the AC domain is sterically hindered by the tight crystal packing around the AC domain (Figure 6B), only the BLUF domain is mobile upon activation, as can also be seen by the overall higher B-factors in the BLUF domain (Supplementary Figure S9). Interestingly, this movement causes a shortening of the unit cell axis a (by 2 Å) and the strong shortening of the b axis (by 5 Å) after light activation compared to the dark state at cryogenic temperature, with a = 52.5 Å, b = 141.4 Å, c = 104.7 Å and a = 54.5 Å, b = 146.4 Å, c = 104.9 Å, respectively.

The final changes observed during signal transmission can be seen in the α-helix linker (α-helix 4). Gln127 to Ser129, as well as the side chain of Phe131 shifting 1 Å closer to the tongue of the AC domain from monomer_B_ (Figure 6D).

Linder *et al*. proposed the tongue between β-sheet 9 and β-sheet 10 to be the link which enables signal transmission from the linker region to the AC domain for bPAC^18^. They observed a structural flexibility in that region, with the tongue being in an *up* position in one monomer and in a *down* position in the other one and suggested this flexibility is of functional relevance for the signal transmission in response to light. Notably in our structure, most of the amino acids in that region also exhibit a high flexibility, as observed by residues in α-helix 4 and the tongue region adopting multiple conformations (Figure 6D).

The changes in our structure do not contradict the idea that the signal is transmitted from the α-helix 3 to the α-helix 4, with a potential transmission to the AC domain via the tongue region. However, as the AC domain is locked in place by several crystal contacts as described above, subsequent changes in the tongue region are not observable with our crystal form.

Finally, some light induced changes are visible in the AC domain of our structure. In particular the C-terminus of OaPAC is significantly disordered in the 5 s cryo-trapped data. Glu326 and Asn327 at the end of β-sheet 13 are the last residues that can be clearly modeled in the electron density of all our modeled structures. Interestingly, their position varies substantially from the dark cryo data, confirming that the C-terminus undergoes a conformational change. As a result, the α-helix 5 seems to be destabilized as well and Asp156 to Pro168 shift slightly. This also explains why the C-terminus of OaPAC has such a strong influence on the cyclase activity^22^, as α-helix 5 is strongly involved in the coordination of the ATP phosphates.

## Conclusion

Our study provides detailed insights into the binding mode of ATP in the active site of native OaPAC and its allosteric influence on the BLUF domain. ATP is mainly coordinated via the phosphate groups and is highly flexible around the adenosine moiety. While at cryogenic temperatures only one conformation is observed, two conformations can be seen at room-temperature. Although, both are a non-productive conformation, it could be shown that this conformation is indeed the native binding mode in OaPAC in the dark. Rotation of the ribose to a productive conformation for cAMP turnover only occurs after light activation and subsequent opening of the active site between both AC domains.

Furthermore, structural information about the early conformational rearrangements in the µs time-scale during activation of the flavin chromophore in the BLUF domain was obtained. Although some of the reported changes in bond distances are at the resolution limit of the crystallographic data, the overall changes in the hydrogen bond network between the dark state and the different light induced states are consistent and therefore trustworthy. The data provide additional information to the ongoing debate about the molecular details during flavin activation in BLUF domains. The cryo-trapped structural data reveal significant conformational changes in the BLUF domain at late reaction times. Our data provide evidence for a Trp_in_/Met_out_ transition upon illumination, not seen in BLUF domains before. We postulate that this switch is involved in signal transmission via β-sheet 5 and the following loop to the linker region (α-helix 3 and 4) and thereby activates the AC domain. However, subsequent movements in the AC domain are hindered by the tight crystal packing. Despite these limitations, our findings contribute to the understanding of how photoreceptor proteins sense and respond to environmental light triggers, with potential implications for the development of optogenetic tools and light-responsive biotechnological applications. Further investigations into the functional implications of these conformational changes will deepen our comprehension of the intricate signaling mechanisms underlying photoactivatable adenylate cyclases.

## Materials and Methods

### Expression and purification of recombinant OaPAC

Sample preparation and injection testing proceeded in the immediate vicinity of the XFEL beamlines in the XBI (XFEL Biology Infrastructure) laboratories^71^.

OaPAC protein used in all experiments corresponds to residues 1 to 350 of the full length OaPAC (UniProt ID K9TLZ5) with C-terminal truncation of 16 amino acids. OaPAC_1-137_ protein used in the FTIR experiments corresponds to residues 1 to 137 of the full length OaPAC (UniProt ID K9TLZ5) with C-terminal truncation of 213 amino acids from the AC domain.

The coding sequence for OaPAC was cloned into a pColdI expression vector (TakaraBio), resulting in pColdI-OaPAC expression vector, encoding for OaPAC_1-350_ in its native form without tag. The coding sequence for OaPAC_1-137_ was cloned into a pColdI expression vector (TakaraBio), the resulting expression vector encoded for OaPAC_1-137_ carrying an N-terminal histidine tag.

pColdI-OaPAC and pColdI-OaPAC_1-137_ expression vectors were transformed into *Escherichia coli* BL21*(DE3) cells, cultured in LB medium and double induced with a cold shock on ice for 5 minutes followed by 10 minutes at 12°C, and the addition of 1 mM Isopropyl β-D-1-thiogalactopyranoside (IPTG) at an OD_600nm_ between 0.4 and 0.6. Cells were grown for an additional 16 h at 30°C. The cells were harvested by centrifugation at 8,000 × g for 30 minutes and the cell pellet was frozen at -20°C until use.

For OaPAC, one cell pellet was resuspended in lysis buffer containing 20 mM Tris/HCl pH 7.5, 20 mM NaCl and protease inhibitors (Roche - cOmplete™ ULTRA Tablets, Mini, EDTA-free, EASYpack Protease Inhibitor Cocktail). After resuspension, cells were lysed by 3 cycles of high-pressure homogenization and centrifuged for 1 h at 30,000 × g. The supernatant was collected and the recombinant protein was purified by fast protein liquid chromatography (FPLC). The first step involved an anion exchange chromatography with elution through a linear salt gradient to 1 M NaCl on a HiTrap Q HP 16/100 column (Cytiva). An ammonium sulfate precipitation was performed with 1.4 M ammonium sulfate. After precipitation, the solution was filtered and the buffer exchanged to 50 mM Tris/HCl pH 8.5, 50 mM NaCl using a PD10 desalting column (Cytiva). The last step consisted of a size exclusion chromatography using a gel filtration column HiLoad 26/600 Superdex 200 pg (Cytiva). Purified protein in 50 mM Tris/HCl pH 8.5, 50 mM NaCl was concentrated to 12 mg/mL, determined by A_280_ measurements using a calculated extinction coefficient of 30160 M^-1^ cm^-1^, flash-cooled in liquid nitrogen and stored at -80°C until use.

For OaPAC_1-137_, one cell pellet was resuspended in lysis buffer containing 10 mM HEPES pH 7.5, 20 mM imidazole, 150 mM NaCl and protease inhibitors (Roche - cOmplete™ ULTRA Tablets, Mini, EDTA-free, EASYpack Protease Inhibitor Cocktail). After resuspension, cells were lysed by 3 cycles of high-pressure homogenization and centrifuged for 1h at 30,000g. The supernatant was collected and the recombinant protein was purified by FPLC. The purification of OaPAC_1-137_ involved an affinity chromatography step on a HisTrap HP column (Cytiva) with elution through a linear imidazole gradient to 500 mM imidazole. After the elution, the protein solution was concentrated and injected onto a gel filtration column HiLoad 26/600 Superdex 200 pg (Cytiva) for size exclusion chromatography. Purified protein in 10 mM HEPES pH 7.5, 150 mM NaCl was concentrated to 12 mg/mL, determined by A_280_ measurements using a calculated extinction coefficient of 28670 M^-1^ cm^-1^, flash-cooled in liquid nitrogen and stored at -80°C until use.

### Protein crystallization and structure determination

Crystals were obtained by vapor diffusion using the sitting drop method. Crystals were grown at 4 °C for one week in the dark after mixing 500 nL OaPAC protein at 10 mg/mL and 500 nL crystallization buffer from the 50 µL reservoir containing 100 mM SPG buffer (succinic acid, sodium dihydrogen phosphate, and glycine in the molar ratios 2:7:7) pH 7.0, 1.2 M disodium succinate, 100 mM guanidine HCl and 5 mM MgCl_2_. Before cryo cooling, a single crystal was soaked with the reservoir solution containing 20 % (v/v) glycerol under red light illumination.

200 × 60 × 40 µm^3^ crystals for dark-state OaPAC grew in an orthorhombic crystal form with space group C222_1_, with one molecule in the asymmetric unit and unit-cell parameters a = 52.9 Å, b = 146.2 Å, c = 103.6 Å, α = 90°, β = 90°, γ = 90°. The high-resolution cutoff was 1.5 Å for this data set. Diffraction data were collected at 100 K at the ID23-2 beamline at ESRF, Grenoble, France using a DECTRIS PILATUS3 X 2M detector^72^. 3,600 frames with 20 ms exposure were recorded with a 360° rotation range and 0.1° oscillation per frame. The X-ray beam was focused to 5 × 26 µm, the energy was set at 14.2 keV and the transmission at 10%.

50 × 35 × 8 µm^3^ crystals for dark-state OaPAC co-crystallized with 5 mM ATP grew in space group C222_1_ with one molecule in the asymmetric unit and unit-cell parameters a = 54.5 Å, b = 146.4 Å, c = 104.9 Å, α = 90°, β = 90°, γ = 90°. The high-resolution cutoff was 2.1 Å for this data set. Diffraction data were collected at 100 K at the ID23-2 beamline at ESRF, Grenoble, France using a DECTRIS PILATUS3 X 2M detector^72^. 3,000 frames with 20 ms exposure were recorded with a 300° rotation range and 0.1° oscillation per frame. The X-ray beam was focused to 5 × 5 µm, the energy was set at 14.2 keV and the transmission at 1%.

For the cryo-trapping experiment, a single 150 × 50 × 30 µm^3^ crystal was illuminated for 5 seconds at 30 mW (10 absorbed photons per molecule) using the illumination system pE-400 (CoolLED) setup at 435 nm. The crystal was flash cooled in liquid nitrogen immediately after illumination. Light-activated OaPAC co-crystallized with 5mM ATP grew in space group C222_1_ with one molecule in the asymmetric unit and unit-cell parameters a = 52.5 Å, b = 141.4 Å, c = 104.7 Å, α = 90°, β = 90°, γ = 90°. The high-resolution cutoff was 1.7 Å for this data set. Diffraction data were collected at 100 K at the P11 beamline at PETRA III, Hamburg, Germany using a DECTRIS EIGER2 X 16M detector^73^. 3,600 frames with 15 ms exposure were recorded with a 360° rotation range and 0.1° per frame. The X-ray beam was focused to 50 × 50 µm, the energy was set at 12.0 keV and the transmission at 35%.

All data were processed and scaled using the program package XDS^74^. All datasets were merged using AIMLESS^75^ from the CCP4 suite^76^. Initial models were obtained by molecular replacement using the PHENIX program suite^77^ with the OaPAC structure from Ohki *et al*. (PDB 4YUS)^20^ as a search model. The models were refined using both the PHENIX program suite^77^ and the CCP4 suite^76^ and manual building was done using COOT^78^. Synchrotron data-structure refinement statistics are shown in supplementary table 1 and structural models have been deposited in the Protein Data Bank under accession codes 8QFE, 8QFE and 8QFG.

All the figures representing the structural models of OaPAC at cryogenic temperature were prepared with the PyMOL Molecular Graphics System, Version 2.0 Schrödinger, LLC^79^.

### Microcrystals growth and TR-SFX experiment

7×3×2 µm^3^ microcrystals were grown in batch after mixing 45% (v/v) OaPAC protein solution at 12 mg/mL, 10 % (v/v) seeds, produced by crushing macrocrystals with seed beads, and 45 % (v/v) crystallization buffer containing 100 mM SPG buffer pH 7.0, 1.4 M disodium succinate, 100 mM guanidine HCl and 5 mM MgCl_2_. Crystals were co-crystallized with 5 mM ATP and grown in the dark at 4°C.

The microcrystal slurry was concentrated to 20% pellet volume (v/v) after sedimentation. The slurry was injected with a gas dynamic virtual nozzle (GDVN)^80^ for a serial femtosecond crystallography (SFX) experiment at the SPB/SFX instrument^81^ of the European XFEL, Schenefeld, Germany in May 2022 (proposal 2829). Data recorded for the experiment at the European XFEL are available at doi:10.22003/XFEL.EU-DATA-002829-00.

OaPAC microcrystals were photoexcited with the second harmonic of the EuXFEL pump—probe laser system^82–84^ at 421 nm. The optical pump pulses were stretched from their original duration of 50 fs to 250 fs by transmission through fused silica. The optical pump had a fluence of 19.8 or 100 mJ/cm^2^ per pulse with a repetition rate of 188 kHz. Under these excitation conditions, 0.6 or 3 photons were absorbed on average per molecule, respectively. The calculation of the absorbed photon number was performed according to the photon calculation Excel sheet by Grübein *et al*.^85^.

The X-ray probe beam was focused to 3 µm × 4 µm, with 9.3 keV photon energy, an estimated pulse length of ≤30 fs and 564 kHz intra-train pulse repetition rate. Data were acquired with the AGIPD detector^86^ operating in fixed medium gain mode. TR-SFX data were collected at 5 ns and 500 ns delays in a LIGHT-DARK1-DARK2 succession pattern (every 3^rd^ pulse within a x-ray pulse train was pumped).

### TR-SFX data processing and dark structure determination

During the experiment, online monitoring was performed with Karabo^87^ and OnDA^88^. Raw data were calibrated using the European XFEL’s internal detector calibration pipeline. Initial hit finding was performed using crystfel 0.10.1^89^ with the following parameters (through the “Extra-xwiz” interface^90^): --peaks=peakfinder8^91^ –min-snr=6 –threshold=200 –min-pix-count=1 –max-pix-count=2 – indexing=none –local-bg-radius=5 –max-res=1200 –min-peaks=0. After manual inspection of the hit/peak finding results, custom python scripts were then used to extract image lists with at least 7 peaks for further processing.

In a second pass, the pre-selected images were subjected to a 2^nd^ round of hit/peak finding with the same parameters as before, apart from –max-pix-count=20, the addition of –int-radius=2,4,6 –min-peaks=10 and no maximum resolution set for peak finding; Indexing was attempted using mosflm^92^, xgandalf^93^, xds^74^ and dirax^94^ in that order.

Images were sorted into “LIGHT”, “DARK1” and “DARK2” based on the pulse IDs; the list of images were annotated accordingly and passed into the partialator program^95^ from CrystFEL for merging using the “—custom-split” option, the unity model, a single iteration and max-ADU set to 110000. All datasets were merged into point group mmm, and the space group was determined to be C222_1_ using AIMLESS^75^ from the CCP4 suite^76^. The high-resolution cutoff was 1.8 Å for all data sets. The space group was C222_1_ and unit-cell parameters a = 54.3, b = 145.8, c = 105.3 Å, α = 90, β = 90, γ = 90°.

The overall crystal hit rate averaged 4.7% from which an average of 85.5% of images could be indexed. However, due to multiple lattices being indexed in some images, the total number of indexed patterns exceeded the total number of hits. A minimum of 40k indexed diffraction patterns were recorded for each data set, and a detailed breakdown of hits and indexed patterns for every dataset are shown in Supplementary Table 2.

The dark structure reported in this manuscript was calculated from all of the DARK2 datasets for the 100 mJ/cm^2^ laser fluence merged together for all time delays. We confirmed that no light contamination was visible in the DARK2 data sets by comparing them to a dark data set collected without optical pump laser. Data collection statistics are shown in Supplementary Table 3. The initial dark model was obtained by molecular replacement using the PHENIX program suite^77^ with the dark-state OaPAC co-crystallized with ATP solved at 100 K as search model.

q-weighted Fourier difference electron density maps were calculated with Xtrapol8^59^ using a fraction of 60,000 diffraction patterns randomly chosen from all DARK2 data collected throughout the beamtime. The dark structure solved from that data set was used to phase the maps. No changes were visible when comparing LIGHT and DARK2 data sets with both laser intensities, showing that the selected time delays (5 ns and 500 ns) are too short to see changes in the electron density. However, the LIGHT-DARK1-DARK2 succession pattern was chosen as light contamination might occur in the first dark pulse data (DARK1), which could be tested for by comparing the DARK1 and DARK2 data. By investigating potentially light contaminated DARK1 data from the 100 mJ/cm^2^ laser fluence exposure, strong differences surrounding the flavin chromophore became visible. This indeed indicates a light contamination of our DARK1 data. Considering the DARK1 data as an additional “light”-state results in an additional time delay of 1.773 µs (X-ray pulse period) between pump and probe pulse. Consequently, this results in a calculated delay of 1778 ns (5 ns plus 1.773 µs) and 2273 ns (500 ns plus 1.773 µs) for the DARK1 data and these new time-delays are referred to as 1.8 µs and 2.3 µs, respectively. A visual representation of the collected pump-probe pattern is shown in supplementary figure S6. It should be noted that the aforementioned absorbed photon number calculation from the pump fluence represents the upper theoretical limit from the LIGHT data and the actual absorbed photons per molecule will be substantially lower in the light-contaminated DARK1 data. Based on this, exact photon dose could not be determined for DARK1 data, but multi-photon effects can be excluded.

After the Fourier difference electron density map calculations between the DARK1 data sets and the DARK2 data set, the occupancy of the light-activated state was determined using Xtrapol8. An occupancy of around 25% was obtained for the two time-delays. Xtrapol8 was also used to calculate extrapolated structure factor amplitudes. These extrapolated measurements generated some negative structure factor amplitudes, but only for less than 5% of structure factors, for each of the two extrapolated data sets (after 1.8 µs and 2.3 µs calculated time delays). The “fref” option^59^ was used in Xtrapol8 in order to estimate their positive values. With this approach, the assumption is that a strong negative difference amplitude is a consequence of measurement errors, and consequently the negative structure factors amplitudes are replaced by their corresponding values in the observed reference data set. This approach will however slightly bias the results towards the reference state, and should therefore only be used when less than 5% of the total structure factor amplitudes are negative. The extrapolated electron density maps obtained after extrapolated structure factors determination were used to model the structural changes happening at the two time points collected.

All TR-SFX models were refined using both the PHENIX program suite^77^ and the CCP4 suite^76^ and manual building was done using COOT^78^. SFX data- and structure refinement statistics are shown in supplementary table 3 and structural models have been deposited in the Protein Data Bank under accession codes 8QFH, 8QFI and 8QFJ. All the figures representing the TR-SFX structural models of OaPAC were prepared with the PyMOL Molecular Graphics System, Version 2.0 Schrödinger, LLC^79^.

### FTIR spectroscopy on OaPAC in solution and in crystal form

Full-length OaPAC and OaPAC_1-137_ proteins in solution were transferred into 10 mM of substrate in 100 mM Tris/HCl, pH 8.5, 50 mM NaCl, 5 mM MgCl_2_ by repeated ultracentrifugation using a Vivaspin 500 filter device with a cutoff of 30 kDa (Sartorius) at 4 °C and afterwards concentrated. The concentration of the samples in crystal form was increased by sedimentation overnight and removal of the supernatant. The final concentration of the samples was 0.3 to 3.0 mM.

1.5 µL of the samples was placed between two BaF_2_ windows, which were sealed without any drying using vacuum grease. The path length was adjusted to reach an absorbance at 1650 cm^-1^ of 0.7 to 1.1. Full hydration was evident from an absorbance ratio of amide I / H_2_O to amide II of 2.7 to 3.5.

The experiments were carried out on an IFS 66/S spectrometer (Bruker) at 10 °C using a mercury cadmium telluride detector. A long wave pass filter with a cut off of 2040 cm^-1^ was used to remove stray light. The spectra were recorded at a scanner velocity of 150 kHz and a spectral resolution of 2 cm^-1^. The Fourier Transform was performed using a zero-filling factor of 4.

The time-resolved difference spectra were obtained using the rapid-scan mode induced by a blue light LED (445 nm, 25 mW/cm^2^, Philips Lumileds) with a diffusion disc placed in front. Full length OaPAC with non-hydrolysable ATP analogue ApCpp, without any substrate, crystallized OaPAC and OaPAC_1-137_ were illuminated for 10 s. Full length OaPAC with ATP was illuminated for 4 s. Data were collected directly before and after the illumination with a sequence of 32, 64, 128, eight times 256 and two times 1024 scans per experiment. For full length OaPAC with ATP, with ApCpp, without any substrate, crystallized OaPAC and OaPAC_1-137_ in total 16, 64, 16, 32 and 32 experiments were averaged, respectively.

For reference measurements, 4 mM of cyclic adenosine monophosphate, adenosine triphosphate, pyrophosphate and phosphate were dissolved in 100 mM Tris/HCl pH 8.5, 50 mM NaCl, 5 mM MgCl_2_ and analyzed using an attenuated total reflection setup. 100 µL of the samples was placed on the internal reflection element with nine active reflections and the buffer was directly subtracted.

### Accession numbers

PDB: 8QFE, 8QFF, 8QFG, 8QFH, 8QFI, 8QFJ

## Author contributions

**Anaïs Chretien**: Investigation, Writing - Original Draft, **Marius Frederik Nagel**: Investigation, Writing - Review & Editing, **Sabine Botha**: Formal analysis, Software, Writing - Review & Editing; **Raphaël de Wijn**: Resources; **Lea Brings**: Resources; **Katerina Dörner**: Resources; **Huijong Han**: Resources; **Jayanath Koliyadu:** Resources; **Romain Letrun**: Resources; **Adam Round**: Resources; **Tokushi Sato**: Resources; **Christina Schmidt**: Resources; **Radu-Costin Secareanu:** Resources; **David von Stetten**: Resources; **Mohammad Vakili**: Resources; **Agnieszka Wrona:** Resources; **Richard Bean**: Resources; **Adrian Mancuso**: Resources; **Joachim Schulz**: Funding acquisition; **Arwen Pearson**: Supervision, Conceptualization; **Tilman Kottke**: Conceptualization, Writing - Review & Editing; **Kristina Lorenzen**: Funding acquisition, Supervision, Conceptualization; **Robin Schubert**: Supervision, Project administration, Funding acquisition, Conceptualization, Writing - Original Draft

## Acknowledgement

S.B. was supported by the NSF Science and Technology Center award 1231306 “Biology with X-ray lasers (BioXFEL)” and the Biodesign Center for Applied Structural Discovery (CASD) at Arizona State University. We acknowledge the European Synchrotron Radiation Facility (ESRF) in Grenoble, France for provision of synchrotron radiation facilities to collect some of the data presented in this research. We would like to thank Dr. Gordon Leonard and Dr. Shibom Basu for assistance and support in using beamline ID23-2. Beamtimes were allocated for proposals MX-2274 and MX-2374. We acknowledge DESY (Hamburg, Germany), a member of the Helmholtz Association HGF, for the provision of experimental facilities. Parts of this research were carried out at PETRA III and we would like to thank Dr. Johanna Hakanpää for assistance in using beamline P11. Beamtime was allocated for proposal BAG-20220737. We acknowledge European XFEL in Schenefeld, Germany, for provision of X-ray free-electron laser beamtime at Scientific Instrument SPB/SFX (Single Particles, Clusters, and Biomolecules and Serial Femtosecond Crystallography) and would like to thank the staff for their assistance. A. R. P. is supported by the Federal Excellence Cluster: CUI: Advanced Imaging of Matter (EXC2056).

