## Supplementary Information for "Light-induced Trpin/Metout switching during BLUF domain activation in ATP-bound photoactivatable adenylate cyclase OaPAC"

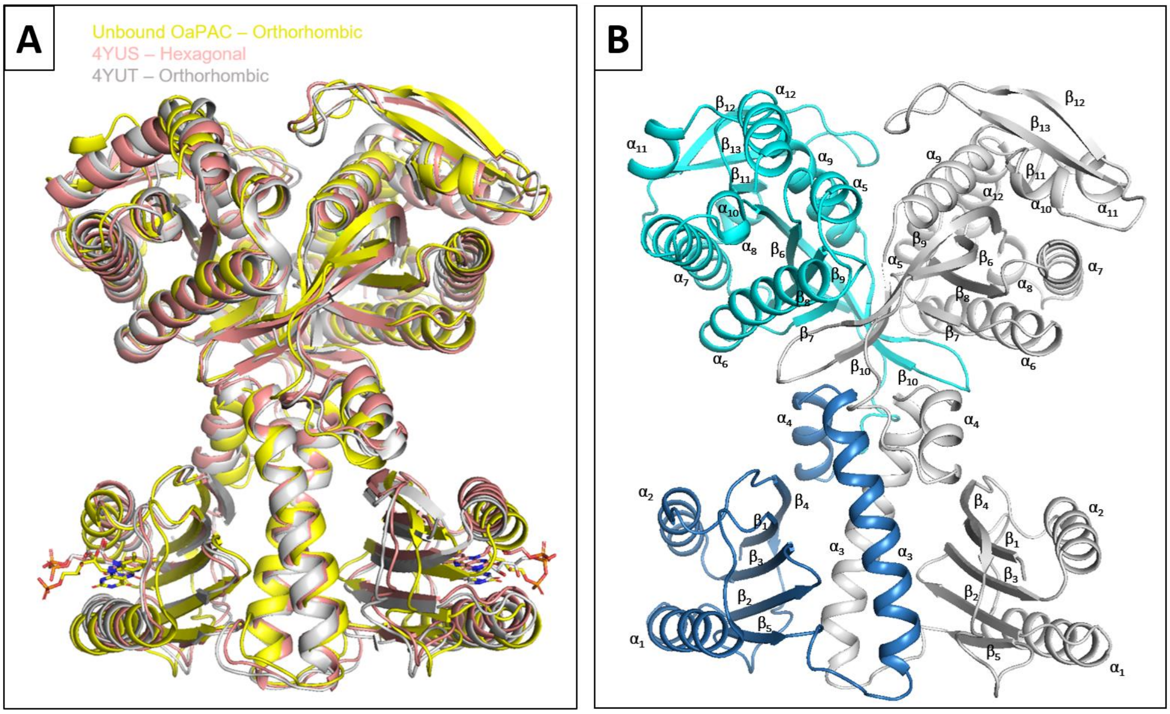


***Figure S 1: Overlay of structural models of OaPAC.*** *A) Model presented in this study in orthorhombic space group (yellow) with hexagonal space group from Ohki et al. (PDB 4YUS)*[1] *(red) and orthorhombic space group from Ohki et al. (PDB 4YUT)*[1] *(grey). B) Annotation of secondary structure elements of OaPAC in cartoon representation with BLUF domain in dark blue and AC domain in cyan color. The overall secondary structure of the ATP-free OaPAC structure determined in this study is comparable to the structures solved by Ohki et al.*[1]*, crystalized either in a hexagonal space group (PDB 4YUS), with 2.19 Å Cα RMSD, or in an orthorhombic space group (PDB 4YUT), with 2.37 Å Cα RMSD. The numbering of the secondary structure elements of OaPAC in this study is according to the labeling shown in (B). Figure prepared by the PyMOL Molecular Graphics System, Version 2.0 Schrödinger, LLC*[2].

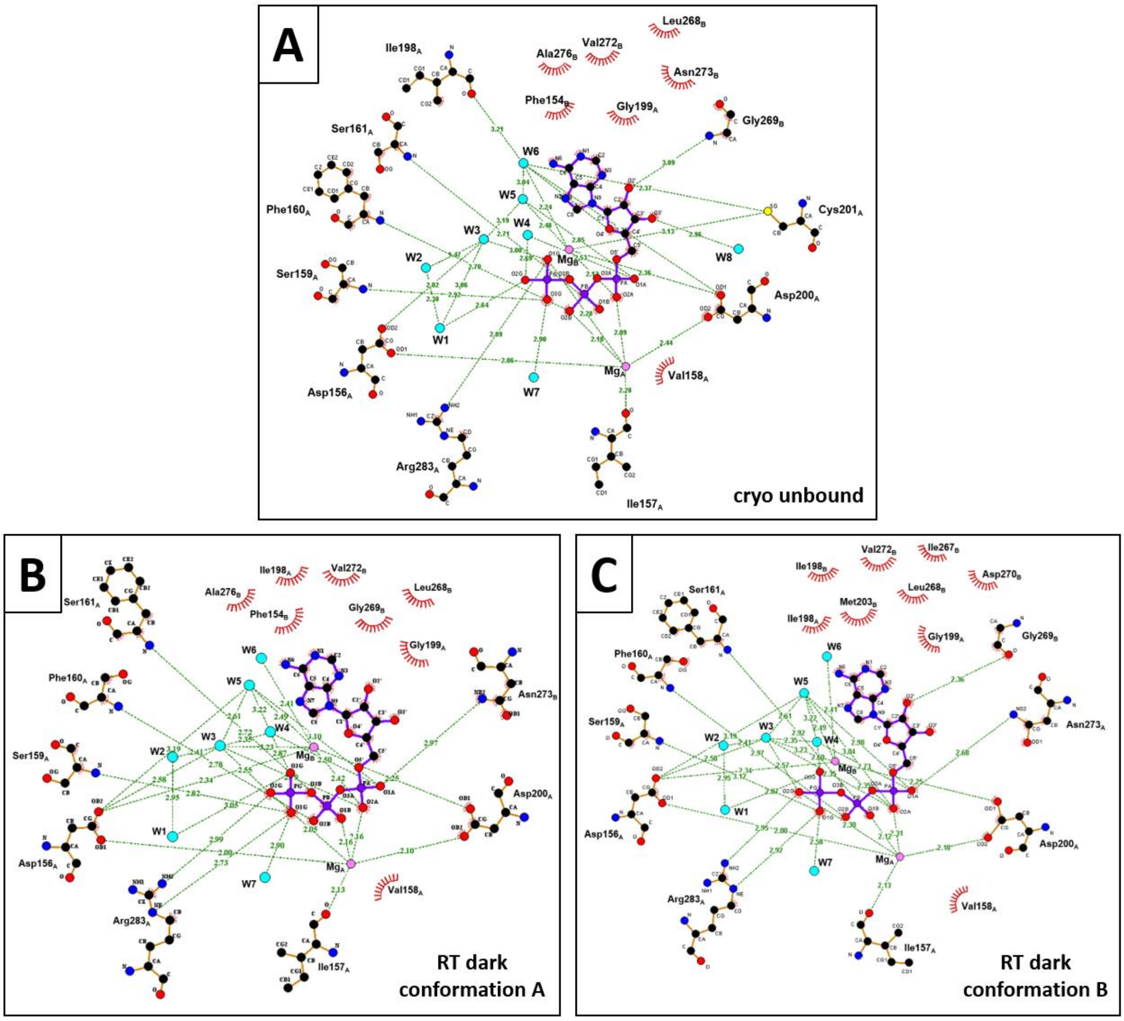


***Figure S 2: Representation of hydrogen bond network coordinating ATP in OaPAC.*** *A) ATP coordination in OaPAC at cryogenic temperature. B and C) ATP coordination in two conformations, both with ~50% occupancy, in OaPAC at room-temperature. As ATP is coordinated mainly via the phosphates, the adenosine moiety has flexibility to adopt both conformations in the active site. While the phosphate group is largely unchanged in both conformations and tightly coordinated, the ribose is rotated 23° in conformation B in respect to conformation A and the adenosine moiety is only loosely coordinated. Figure prepared by LigPlot$^{+}$ v2.2*[3].

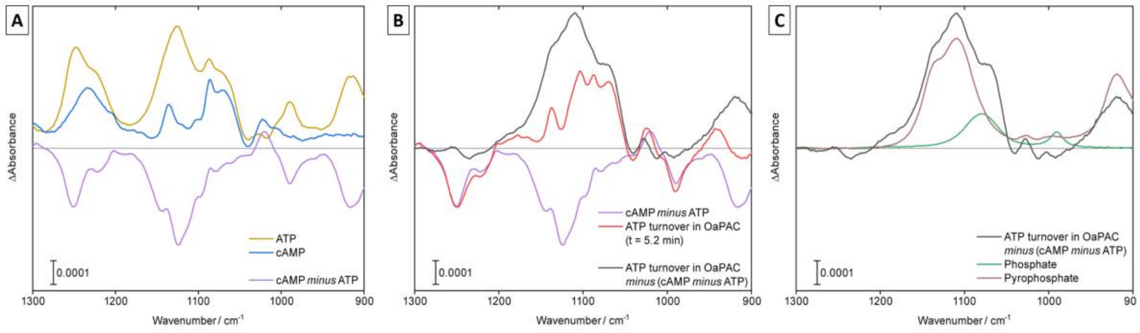


***Figure S 3: FTIR light-minus-dark difference spectra of OaPAC in the spectral region, which shows features of conversion from ATP to cAMP.*** *A) Reference spectra of ATP and cAMP in solution are shown and result in the difference spectrum 'cAMP minus ATP'. B) Subtraction of the difference spectrum 'cAMP minus ATP' (shown in A) from the difference spectrum of OaPAC in solution at 5 min after illumination isolates the contributions of byproducts of the conversion from ATP to cAMP C) The resulting spectrum from (B) matches well the reference spectra of contributions from phosphate and pyrophosphate in solution, which are the byproducts during ATP hydrolysis to cAMP. Concluding, the difference signals of OaPAC after illumination can be clearly assigned to ATP and the products of its photo-induced turnover to cAMP.*

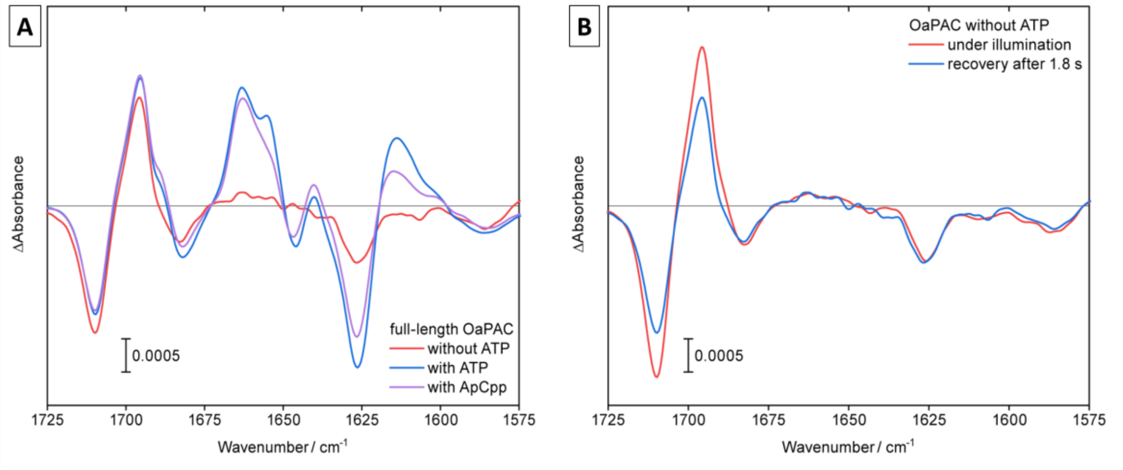


***Figure S 4: Presence of ATP and the non-hydrolysable analog ApCpp strongly influences the light-minus-dark FTIR difference spectrum of OaPAC.*** *A) In the absence of ATP, the contributions from the AC domain of OaPAC to the difference spectrum nearly diminish completely. Addition of ApCpp leads to a partial recovery of the light-induced response of the AC domain. B) No recovery is observed after light exposure from the region attributed to AC contribution, while relaxation in the peaks from the BLUF domain is observed (1994 and 1710 cm$^{-1}$). Accordingly, the bands of the AC response are nearly completely suppressed without any substrate bound. Even under illumination, any significant signals are not detectable.*

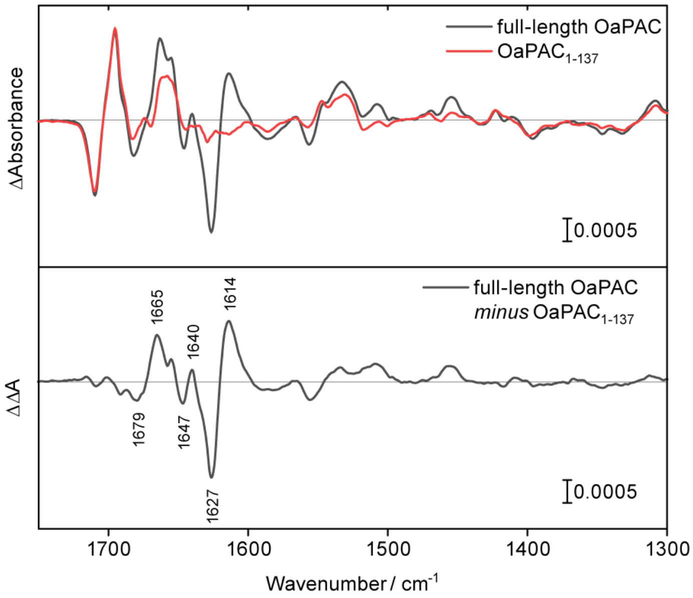


***Figure S 5: Light-minus-dark FTIR difference spectra of full-length OaPAC and OaPAC$_{1\text{-}137}$ and their double difference spectrum isolating contributions by the presence of the AC domain**. To reveal which domain contributes to which bands, a truncated construct containing only the BLUF domain and linking helices $\alpha$3 and $\alpha$4 (OaPAC1-137) was generated (upper panel). An overlay with the difference spectrum of OaPAC reveals that most of the prominent bands between 1680 and 1600 cm$^{-1}$ originate from the presence of the AC domain but for one signal at 1655 cm$^{-1}$ which is still present in the truncated construct.*

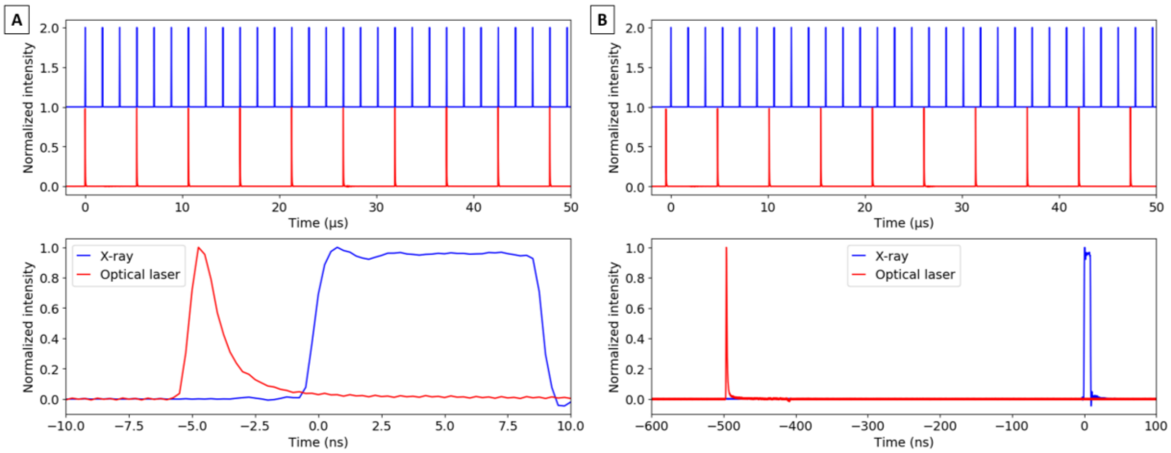


***Figure S 6: Oscilloscope trace of the TR-SFX experiment illustrating the applied pump-probe pulse pattern in LIGHT-DARK1-DARK2 succession****. A) Data were recorded with a time delay between pump pulse (red) and first probe pulse (blue) of 5 ns (LIGHT) and 1778 ns for the second probe pulse (DARK1). Note that the presented probe pulse is used for laser synchronization and does not represent the actual x-ray pulse. The actual x-ray pulse is timed in the rising slope of the pump pulse. B) Data was recorded with a time delay between pump pulse (red) and first probe pulse (blue) of 500 ns (LIGHT) and 2273 ns for the second probe pulse (DARK1).*

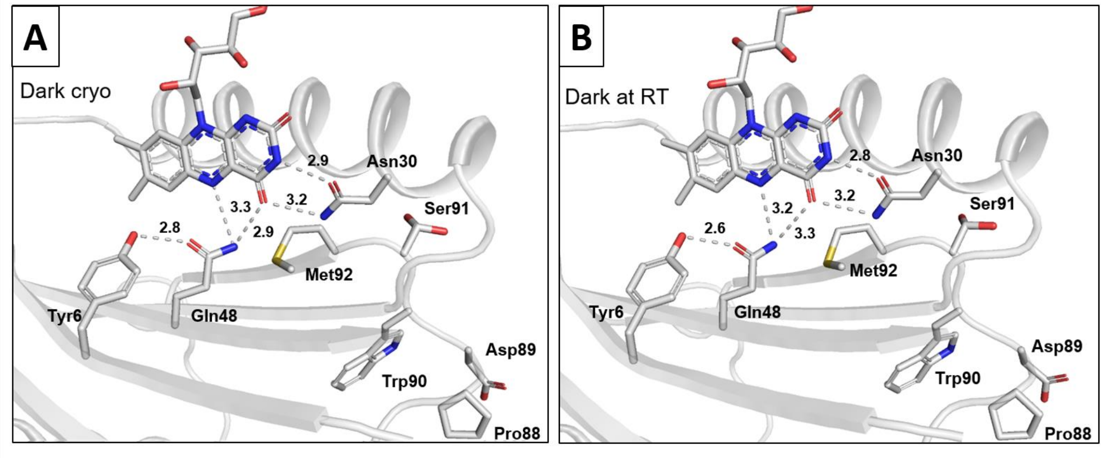


***Figure S 7: Hydrogen bond network around FAD chromophore in dark state OaPAC at cryogenic temperature (A) and room-temperature (B).*** *Figure prepared by the PyMOL Molecular Graphics System, Version 2.0 Schrödinger, LLC[2].*

.

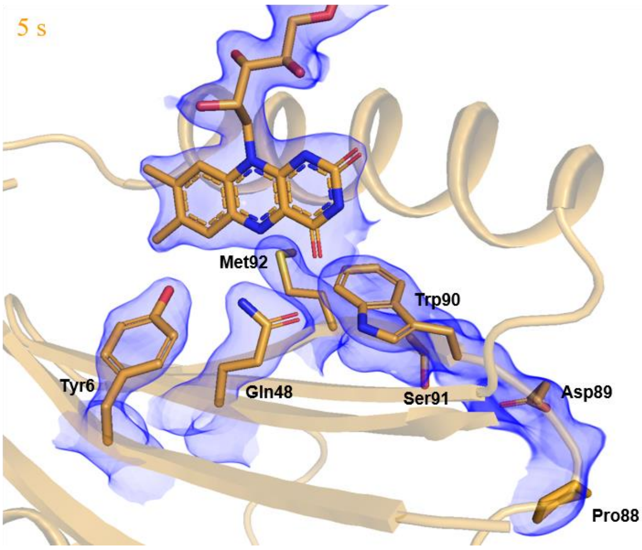


***Figure S 8: Electron density map ($2F_o$-$F_c$) contoured at 1 σ around FAD chromophore in 5s cryo-trapped data from OaPAC showing the Trp-90$_{in}$/Met-92$_{out}$ conformation.*** *Figure prepared by the PyMOL Molecular Graphics System, Version 2.0 Schrödinger, LLC[2].*

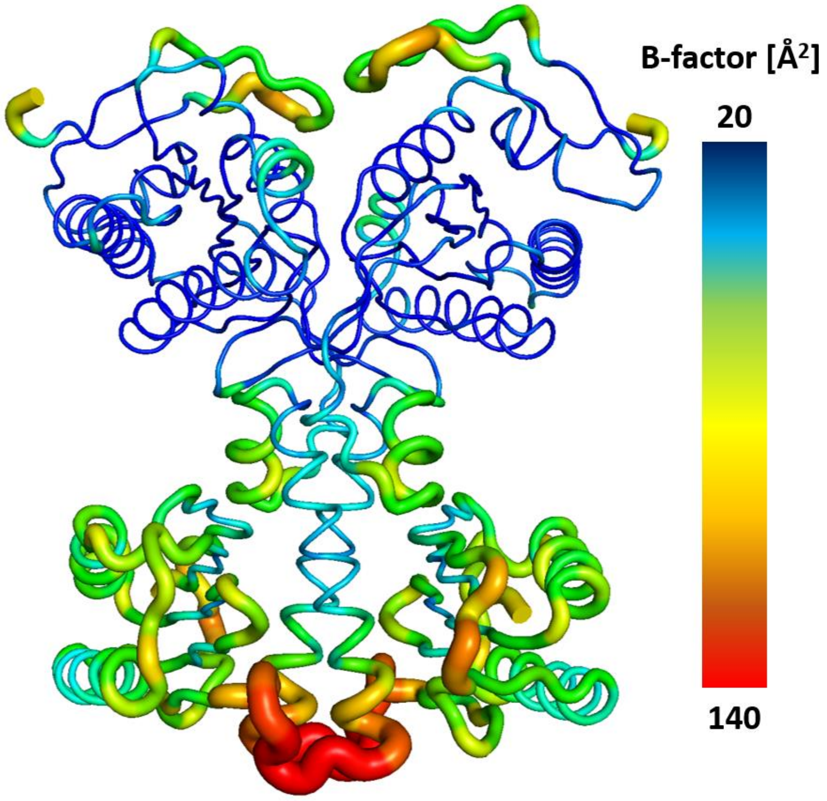


***Figure S 9: OaPAC secondary structure from 5 s cryo trapped data in B-factor putty presentation.*** *Lower temperature factors colored in blue indicate good agreement of the data with the structural model and low flexibility, while high temperature factors colored in red indicate a high degree of flexibility, indicated also by the thicker ribbon presentation. Figure prepared by the PyMOL Molecular Graphics System, Version 2.0 Schrödinger, LLC[2].*

***Supplementary video 1:*** *Interpolation of the amino acid side chain movements and conformational changes in the surrounding of the chromophore in the BLUF domain of OaPAC. Starting model is from the dark RT-SFX structure, intermediate models are from the 1.8 µs and 2.3 µs TR-SFX structures and final model is from the 5 s cryo-trapped structure. Video is prepared by the PyMOL Molecular Graphics System, Version 2.0 Schrödinger, LLC[2] using the "morph" command and linear model for interpolation.*

*Supplementary table 1: Synchrotron data collection and processing statistics*

| OaPAC | Cryo ATP-free | Cryo ATP bound | 5 s illuminated cryo-trapped |
|---|---|---|---|
| **PDB deposition code** | 8QFE | 8QFF | 8QFG |
| **Data collection** | | | |
| Source | ESRF (ID23-2) | ESRF (ID23-2) | PETRA III (P11) |
| Temperature (K) | | 100 | |
| Wavelength (Å) | 0.8731 | 0.8731 | 1.0332 |
| Detector | DECTRIS PILATUS3 X 2M | DECTRIS PILATUS3 X 2M | DECTRIS EIGER2 X 16M |
| Space group | | $C222_1$ | |
| Cell dimensions | | | |
| *a*, *b*, *c* (Å) | 52.9, 146.2, 103.6 | 54.5, 146.4, 104.9 | 52.5, 141.4, 104.7 |
| α, β, γ (°) | 90, 90, 90 | 90, 90, 90 | 90, 90, 90 |
| Solvent Content (%) | 51.5 | 53.7 | 50.2 |
| Resolution (Å) | 33.86-1.50 (1.53-1.50) | 60.04-2.10 (2-16-2.10) | 42.08-1.70 (1.73-1.70) |
| $R_{meas}$ (%) | 5.5 (131.2) | 23.0 (211.5) | 3.6 (94.4) |
| <*I*> / sig*I* | 19.7 (1.1) | 9.0 (1.2) | 34.9 (2.7) |
| $CC_{1/2}$ | 1.000 (0.846) | 0.997 (0.501) | 1.000 (0.939) |
| No. unique reflections | 64,596 (3,212) | 24,956 (2,002) | 42,861 (2,140) |
| Completeness (%) | 100.0 (100.0) | 100.0 (100.0) | 99.2 (95.7) |
| Redundancy | 13.0 (12.8) | 10.9 (11.3) | 13.3 (12.0) |
| **Refinement** | | | |
| Resolution (Å) | 33.22-1.50 (1.52-1.50) | 60.04-2.10 (2.20-2.10) | 42.08-1.70 (1.74-1.70) |
| Completeness (%) | 99.96 (99.96) | 99.96 (100.00) | 98.63 (95.33) |
| No. unique reflections | 64,543 (2,636) | 24,924 (2,935) | 42,714 (2,559) |
| $R_{work}$ / $R_{free}$ | 0.163 / 0.186 | 0.189 / 0.223 | 0.209 / 0.221 |
| No. atoms | | | |
| Protein | 3,097 | 2,979 | 2,608 |
| Ligand/ion | 50 | 93 | 93 |
| Water | 328 | 189 | 107 |
| *B*-factors | | | |
| Protein (Å$^2$) | 30.44 | 45.33 | 60.14 |
| Ligand/ion (Å$^2$) | 24.53 | 46.05 | 49.86 |
| Water (Å$^2$) | 40.150 | 47.55 | 49.88 |
| r.m.s. deviations | | | |
| Bond lengths (Å) | 0.013 | 0.010 | 0.014 |
| Bond angles (°) | 1.190 | 1.020 | 1.160 |
| Ramachandran | | | |
| favored (%) | 96.55 | 97.70 | 93.88 |
| allowed (%) | 3.16 | 2.01 | 6.12 |
| outliers (%) | 0.29 | 0.29 | 0 |
| Clashscore | 3.34 | 2.12 | 7.13 |

*Values in parenthesis for highest resolution shell.*

*Supplementary table 2: XFEL hit statistics for all datasets*

| Pump delay [ns] | Laser Fluence [mJ cm$^2$] | # Frames | Hits | Indexed Lattices | | | |
|---|---|---|---|---|---|---|---|
| | | | | Total | Light | Dark1 | Dark2 |
| 5 (1.8 µs) | 100 | 3,771,564 | 147,201 | 153,247 | 51,838 | 50,786 | 51,247 |
| 500 (2.3 µs) | 100 | 3,144,645 | 205,695 | 210,575 | 70,798 | 68,993 | 70,784 |

### *Supplementary table 3: Data collection and refinement statistics of SFX data.*

| OaPAC | Dark-RT (all frames) | Dark (60k subset) | 1.8 µs delay | 2.3 µs delay |
|---|---|---|---|---|
| **PDB deposition code** | 8QFH | - | 8QFI | 8QFJ |
| **Data collection** | | | | |
| Source | EuXFEL (SPB/SFX instrument) | | | |
| Temperature (K) | 293 | | | |
| Wavelength (Å) | 1.3332 | | | |
| Detector | AGIPD | | | |
| Space group | C2221 | | | |
| Cell dimensions | | | | |
| a, b, c (Å) | 54.3, 145.8, 105.3 | | | |
| α, β, γ (°) | 90, 90, 90 | | | |
| Solvent Content (%) | 53.6 | | | |
| Resolution (Å) | 31.63-1.80 (1.86-1.80) | 31.63-1.82 (1.88-1.82) | 31.63-1.82 (1.88-1.82) | 31.63-1.82 (1.88-1.82) |
| # patterns merged | 299,140 | 59,892 | 50,786 | 68,993 |
| $R_{split}$ (%) | 5.4 (41.6) | 10.5 (74.1) | 13.2 (171.1) | 11.0 (157.0) |
| <*I*> / sig*I* | 11.6 (2.6) | 6.2 (1.5) | 1.2 (0.6) | 1.3 (0.8) |
| $CC_{1/2}$ | 0.997 (0.876) | 0.987 (0.645) | 0.983 (0.391) | 0.988 (0.448) |
| CC* | 0.999 (0.996) | 0.997 (0.886) | 0.996 (0.750) | 0.997 (0.787) |
| No. unique reflections | 39,352 (3,858) | 39,352 (3,858) | 36,898 (3,192) | 37,632 (3,250) |
| Completeness (%) | 100.00 (100.00) | 100.00 (100.00) | 96.61 (84.92) | 98.52 (86.37) |
| Redundancy | 3,079 (1,836) | 448 (267) | 711 (428) | 936 (563) |
| **Refinement** | | | | |
| Resolution (Å) | 31.63-1.80 (1.82-1.80) | | 31.63-1.90 (1.96-1.90) | 31.63-1.90 (1.96-1.90) |
| Completeness (%) | 99.80 (99.93) | | 99.92 (99.85) | 99.89 (99.89) |
| No. unique reflections | 39,103 (12,73) | | 33,355 (2,609) | 33,344 (2,609) |
| $R_{work}$ / $R_{free}$ | 0.165 / 0.194 | | 0.307/ 0.340 | 0.294 / 0.354 |
| No. atoms | | | | |
| Protein | 2,988 | | 2,782 | 2,770 |
| Ligand/ion | 135 | | 135 | 135 |
| Water | 81 | | 129 | 127 |
| *B*-factors | | | | |
| Protein (Å$^2$) | 59.02 | | 41.89 | 46.12 |
| Ligand/ion (Å$^2$) | 44.47 | | 34.19 | 43.73 |
| Water (Å$^2$) | 56.91 | | 41.64 | 42.17 |
| r.m.s. deviations | | | | |
| Bond lengths (Å) | 0.007 | | 0.014 | 0.006 |
| Bond angles (°) | 0.93 | | 0.95 | 0.81 |
| Ramachandran | | | | |
| favored (%) | 97.13 | | 97.26 | 96.04 |
| allowed (%) | 2.59 | | 2.74 | 3.96 |
| outliers (%) | 0.29 | | 0 | 0 |
| Clashscore | 3.88 | | 4.30 | 5.38 |

*Values in parenthesis for highest resolution shell.*
